# Characterising sleep spindle emergence in infancy: Effects of prematurity and early locomotor intervention

**DOI:** 10.64898/2026.10.20.760527

**Authors:** Amaia Dornier, Parvaneh Adibpour, Valérie Biran, Amandine Pedoux, Hala Nasser, Adrien Conessa, Aline Gonzalez Carpinteiro, Pauliina Yrjölä, Laurie Devisscher, Léa Guéret, Elodie Hinnekens, Alice Heneau, Olivier Sibony, Lucie Hertz Pannier, Marianne Alison, Richard Delorme, Marianne Barbu Roth, Jessica Dubois, Arnaud Boutin

## Abstract

**Study objectives:** Sleep spindles are thalamocortical oscillations evolving during early brain maturation. In adults, they support memory consolidation and are organised into rhythmic “trains” thought to support memory reactivation. How this temporal organisation emerges in infancy, and whether prematurity or early sensorimotor experience shapes it, remains unclear. This study characterised infant spindles and investigated how prematurity and early locomotor training affect their temporal and spatial organisation, providing insight into early brain maturation.

**Methods:** Daytime high-density EEG sleep recordings were analysed in 50 infants born full-term (n=12) or very preterm (n=38, including 12 who received locomotor training) at 2 months corrected age. Spindle temporal organisation was analysed using a two-step Morlet wavelet transform to quantify slow modulations of spindle-band power (11-16Hz). Time–frequency decomposition around spindle onset was used to identify significant spindle-band scalp-wide activity, and topographical maps were statistically compared between groups.

**Results:** All groups showed rhythmic spindle organisation, three times slower than in adults, with no group differences. Untrained preterm infants showed a more diffuse spindle-band activity than the distinct frontal and central clusters observed in full-term infants. Trained preterm infants exhibited a spatial distribution closely resembling that of full-term infants, although their frontal spindle density was lower than in both other groups.

**Conclusions:** Spindle rhythmic organisation emerged early in infancy and remained unaffected by prematurity or locomotor training. Prematurity was associated with more diffuse spindle-band activity, suggesting reduced cortical specialisation, whereas trained preterm infants showed a spatial distribution resembling that of full-term infants, suggesting that early locomotor experience may influence thalamocortical organisation.

**Statement of significance:** This study examined sleep spindles at the age of their emergence, as valuable markers of brain development linked to thalamo-cortical networks and maturation. First, we showed that their rhythmic organisation and central/frontal spatial distribution are already observed in infants from 2 months of age, suggesting that early processes of memory consolidation are at play in relation to sensorimotor development. We also observed that early life factors such as prematurity and locomotor training modulate spindle characteristics. These findings make spindles relevant early markers for identifying infants at risk of atypical developmental and for assessing the effects of interventions capitalising on the brain’s early plasticity. Future longitudinal studies will be needed to determine whether these early cerebral observations yield lasting developmental benefits.

## Introduction

Sleep plays a crucial role in early brain development by driving the formation of neural networks ([1], see [2,3] for reviews) and supporting offline memory consolidation (e.g. [3,4], see [5] for a review). The developing sensorimotor system particularly relies on this process to establish its underlying thalamo-cortical circuits ([6], see [2,3] for reviews). As non-rapid eye movement (NREM) sleep develops, changes in sleep spindle characteristics may serve as indicators of brain maturation. These neural oscillations, generated through interactions between the thalamus and the cortex, can be recorded via scalp electroencephalography (EEG) and are widely considered key markers of thalamo-cortical network development [7–9]. However, despite the established importance of sleep spindles in development, little is known about their temporal organisation in early infancy nor how they are modulated by early life factors.

In adults, sleep spindles are characterised by a duration of approximately 0.3–2 seconds (s) and a frequency range of 11–16 Hz. They are known to be implicated in sleep-dependent memory consolidation (e.g. [4], see [3,5] for reviews focusing on procedural memory consolidation). Recent studies have shown that sleep spindles do not appear randomly over time, but instead follow a structured rhythmic pattern that is closely linked to the efficiency of memory reactivation and consolidation. Rather than occurring only as isolated events, spindles can also cluster into transient spindle-rich periods, often referred to as spindle trains (see [5] for a review). This temporal cluster-based organisation involves two nested rhythms: an infraslow rhythm of approximately 0.02 Hz, corresponding to the interval between successive spindle trains, and a mesoscale rhythm of approximately 0.2-0.3 Hz, corresponding to the interval between two spindles within a train. This fine alternation between spindle events and refractory periods is thought to support the temporal separation of memory reactivation episodes, thereby limiting interference between competing memory traces ([10], see [5] for a review). Therefore, investigating this temporal organisation in infants serves a dual purpose: revealing how memory reactivation episodes are structured during early development, and reflecting the maturation state of the underlying thalamo-cortical network.

In infants, sleep spindles are not present at birth but typically emerge during early postnatal life, at around 3 to 9 weeks of age [9]. Upon their emergence, spindles show a predominantly frontopolar and central spatial distribution. Their duration is generally longer in infants than in adults: it increases between approximately 1 and 4 months of age (M), then it gradually declines over the first postnatal year [9]. In parallel, spindle density increases almost linearly over the first months to reach adult values while spindle frequency remains relatively stable within the 11–16 Hz range across early development. However, the development of spindle rhythmicity (i.e., infraslow and mesoscale rhythms) remains largely unexplored despite its established critical role in the functional efficiency of the thalamo-cortical network ([10], see [5] for a review). To date, only a single exploratory MEG study has investigated spindle rhythmicity in a small sample of seven infants across a broad age range (46 to 63 weeks GA) providing preliminary indications of a potential rhythmic pattern [11]. Consequently, we aimed to determine whether sleep spindles, at their emergence in infancy, already exhibit a temporal organisation similar to that observed in adults, or whether this temporal organisation emerges progressively over early brain maturation.

Furthermore, through their role in early thalamo-cortical development and procedural memory consolidation, infant sleep spindles may also contribute to sensorimotor plasticity. At 3M, some synchrony emerges between subcortically generated limb twitches and sleep spindles localised over sensorimotor cortical regions associated with the moving limb [12]. This coordinated activity provides the sensorimotor system with sensory feedback that enriches cortical sensorimotor maps, driving motor learning and memory via the cortico-hippocampal and thalamo-cortical networks ([6], see [3]for a review). Furthermore, in 6M-old infants, [13] highlighted that the spatial topography of fast spindle density serves as an indicator of sensorimotor development. Increased density localised in the central brain regions was associated with higher gross motor scores at 12M, whereas a more widespread frontocentral density was associated with improved gross motor scores at 24M. Consequently, characterising sleep spindles—and their spatial distribution in particular— represents particularly promising markers for assessing the quality of early developmental trajectories of the sensorimotor system and also for identifying potential risks of neurodevelopmental disorders (see [14] for a review).

Indeed, several studies have suggested that sleep spindle characteristics can be altered in atypical developmental conditions (see [14,15] for reviews). The formation and maturation of thalamocortical connections takes place during the second half of pregnancy [16], a developmental window during which these circuits appear particularly vulnerable to perturbations such as preterm birth. Consequently, preterm born infants provide a particularly relevant population for investigating how early alterations in brain maturation may affect the emergence and organisation of sleep spindles. Numerous studies have reported developmental impairments in preterm born infants at both behavioural (e.g., [17]) and cerebral levels (e.g., in EEG: [18–20] / in MRI: [21], see [22] for a review) particularly in the sensorimotor domain and thalamo-cortical connections (e.g., [23–26]). Considering the growing evidence that sleep spindles reflect the maturation of thalamo-cortical circuits, including sensorimotor pathways, prematurity may be expected to influence sleep spindle development, but this remains largely understudied.

Importantly, although adverse perinatal conditions such as prematurity may be associated with altered early developmental trajectories, infancy is a period of intense brain plasticity marked by rapid structural reorganization and dendritic growth [27] allowing experience-dependent modulation of trajectories. Early interventions might then improve neurodevelopmental outcomes ([28,29], see [30,31] for reviews). However, their impact on sleep spindle characteristics remain to be tested, particularly when considering sensorimotor interventions given their evidenced role in sensorimotor plasticity ([6,12], see [2,3] for reviews). This is an intriguing question raised by recent studies proposing an early locomotor training which improves outcome in preterm born infants [32]. It is based on a mini-skateboard adapted for newborns and young infants (the *Crawliskate*), and designed to facilitate early quadrupedal locomotion by supporting comfortable prone positioning, elevating the head and trunk, and allowing free and coordinated movement of the arms and legs necessary for self-propelled crawling. Influenced by sensory inputs such as visual cues, the mother smell or tongue [33–35], this early locomotion is not merely a reflex but is thought to involve the brain’s sensorimotor networks. Nevertheless, the brain substrates of behavioural benefits observed in preterm infants [32] remain to be explored, and sleep spindles may represent a promising candidate marker given their reported role in the development of sensorimotor networks.

In the present study, we thus aimed to determine whether sleep spindles in early infancy already display a structured temporal organisation resembling that observed in adults, whether their temporal organisation and spatial distribution are altered by prematurity, and whether this might be modulated by early locomotor training. We recorded EEG during a daytime nap in full-term infants at 2M and in very preterm infants assessed at 2M of corrected age (CA). A subcohort of preterm infants underwent an 8-week Crawliskate training at home (between term-equivalent age and 2M CA). We hypothesised that sleep spindles at 2M would already exhibit a structured, rhythmic temporal organisation in all infants, with a spatial distribution involving mainly central and frontal regions. Further, we hypothesised that prematurity would alter sleep spindle characteristics, with more immature and irregular temporal dynamics, and less pronounced spatial clustering in central and frontal regions. Finally, we expected the Crawliskate training to partially compensate for the developmental alterations associated with prematurity, leading to a pattern more similar to the one observed in full-term infants.

## Method

The study was conducted in accordance with the ethical principles of the Declaration of Helsinki. The design and procedures were reviewed and approved by the ethics committee (Comité de Protection des Personnes, CPP Ile de France 3) and by the ANSM (Agence Nationale de Sécurité du Médicament et des produits de santé). After receiving detailed information about the study, the parents provided written informed consent before their infant’s participation.

### Participants

#### Full-term infants

A typically-developing control group of 17 full-term infants was included in the study (mean of gestational age (GA) at birth: 40.30 +/- 1.10 weeks, range: 37.60-41.60w, 9 girls). They had no neurological risk or background (i.e., incident-free pregnancy and birth, no postnatal complications, and no immediate family members with neurological or genetic abnormalities). To ensure infants met these criteria, we relied on the guidelines established by the French “Haute Autorité de Santé” regarding risk factors for neurodevelopmental disorders [36]. Among those 17 infants, 12 had EEG recordings meeting the inclusion criteria as described above and were included in the following analyses (mean of GA : 40.20 +/- 1.30w, range : 37.60-41.60w, 7 girls).

#### Preterm infants

This study also included 52 preterm infants (24 girls) born before 33w GA (mean GA at birth: 2.6 +/- 2.3w, range: 24.1–32.9w. Among these infants, 38 met the EEG inclusion criteria described below and were included in the subsequent analyses (mean GA at birth: 27.7 ± 2.2w, range: 24.1–32.4w, 21 girls). Based on their GA at birth, infants were categorised into three groups : G1 (<26w), G2 (26w-27w + 6 days), G3 (≥28w) (see table 1).

**Table 1:**
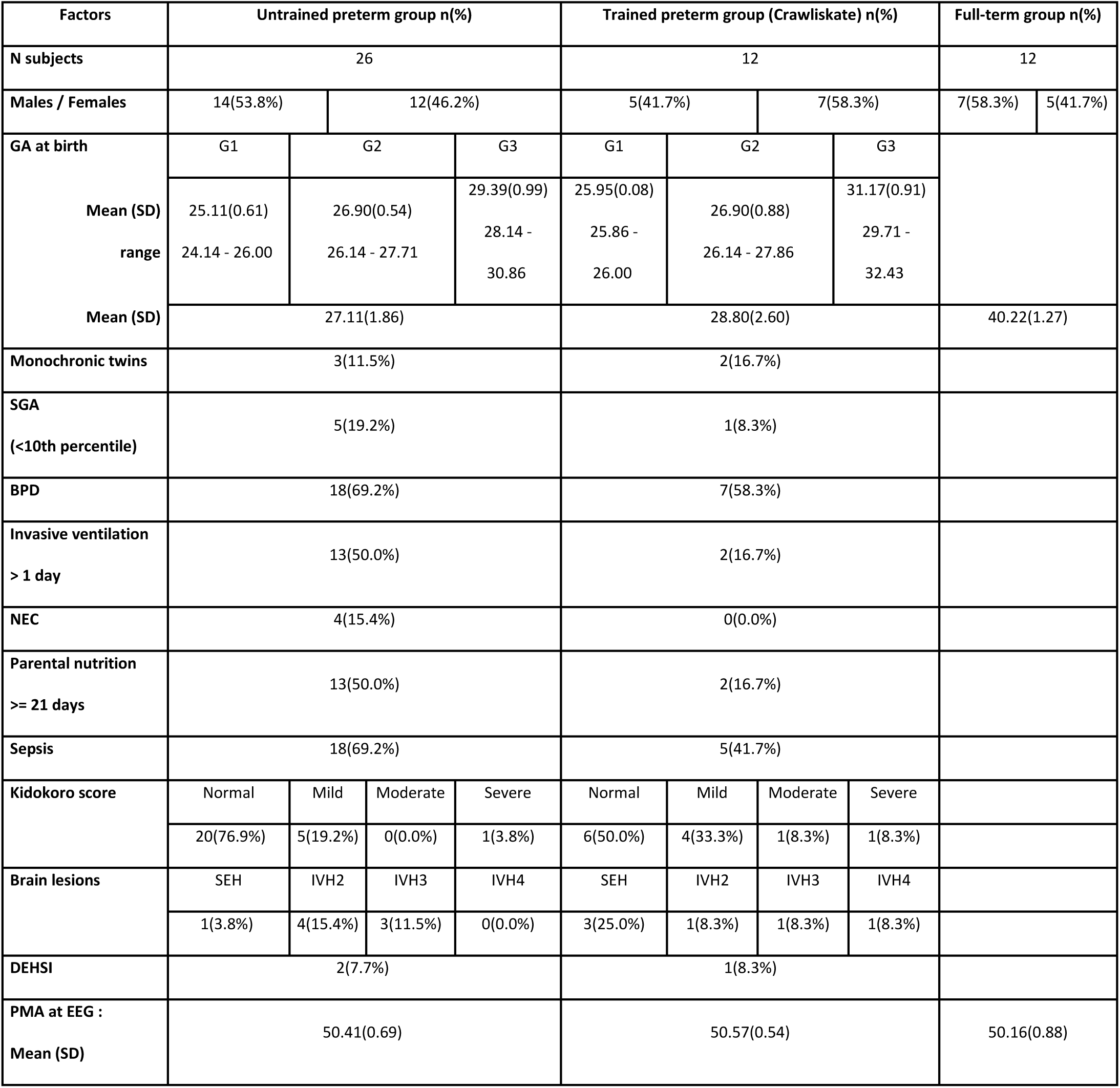
Characteristics of the cohort of full-term and preterm infants. GA: Gestational Age ; SD: Standard Deviation ; SGA: Small for Gestational Age ; BPD: Bronchopulmonary Dysplasia ; NEC: Necrotizing Enterocolitis ; SEH: Subependymal Hemorrhage ; IVH: Intraventricular Hemorrhage; DEHSI: Diffuse Excessive High Signal Intensity.

A set of relevant clinical risk factors were recorded systematically [37,38]. These included monochronic twin pregnancy, being small for GA at birth (SGA, <10^th^ percentile), the occurrence of bronchopulmonary dysplasia (BPD), the use of invasive ventilation (>1 day), the presence of necrotising enterocolitis (NEC), the use of prolonged parental nutrition (≥21 days), and perinatal sepsis. In addition to EEG examinations (detailed below), these infants were due to have a clinical MRI exam at TEA because of GA lower than 28w or clinical/neurological suspicions. This exam allowed us to measure the Kidokoro score [39], which provides overall information on brain maturation and injury, and consequently on the risk of later neurodevelopmental impairment [40] (see table1).

##### Locomotor training

Among the 52 preterm infants included in the study, 19 were randomly assigned to the training group, leading to a group of 12 trained infants over the 38 included in the analyses (31.6%) (see Table 1 for details on the characteristics of each preterm infant group). Upon NICU discharge, they followed an 8-weeks home training intervention on the Crawliskate, consisting of daily 10-minute parent-led and professional-led (physical therapists) sessions (average training time over 8 weeks: 357.99 min +/- 107.83, range = 182.33 – 493.93). Training started when infants had reached 37–42w of GA and had been discharged from the NICU. It continued until 2M CA, prior to EEG acquisition. Detailed information on the training is provided in [32] and in Supplementary information, Table 1.

To ensure that the groups of trained and untrained preterm infants were comparable, Mann–Whitney tests were conducted to determine whether GA at birth and age at the time of EEG acquisition differed significantly. No significant difference was found in terms of GA at birth (U = 99, *p* = 0.08), nor in age at EEG acquisition (U = 115, *p* = 0.20). To ensure that our findings were not driven by gestational age at birth or group heterogeneity regarding Kidokoro scores and brain abnormalities, we conducted supplementary analyses. Specifically, we selected a subgroup of untrained preterm infants whose age and brain abnormality profiles closely matched those of the trained infants. We confirmed that the results remained consistent within this matched subset (see Supplementary Information, Table 2, Figure 1).

#### EEG examinations

All full-term and preterm infants underwent high-density EEG recordings for research purposes using a 128-channel high density (HD)-EEG system (Magstim EGI, Eugene, USA, 128-channels), with a sampling rate of 1000 Hz. EEG acquisition was performed at 2M of age for full-term infants (mean of PMA: 50.20 +/- 0.90w, range: 48. 90-51.90w) and at 2M CA for preterm infants (mean of PMA: 50.46 +/- 0.65w, range: 49.10-52.00w) (Table 1). Acquisitions were carried out during the infant’s spontaneous sleep during a daily nap for approximately 10 minutes. During acquisition, infants rested in a baby cot, or occasionally in the parent’s arms when this helped them remain calm [18]. A video camera was used to record the infant’s behaviour and the experimenter manually marked approximate vigilance states based on eye openings and movements. These facilitated subsequent offline sleep scoring by an expert.

### Sleep scoring

To assess vigilance states, EEG data were first transformed into a low-density bipolar montage, as commonly used in standard clinical settings. The recordings were then visually inspected and scored into different vigilance states by an expert in neonatal electrophysiology (H.N.) (see [18], for details). Vigilance states were categorised as wakefulness, rapid eye movement (REM) sleep, and non-rapid eye movement (NREM) sleep. The different states were distinguished from one another using the same criteria provided by [41]. REM sleep was characterised by irregular, low-voltage, or mixed activity patterns, whereas NREM sleep was identified by the emergence of sleep spindles. Wakefulness was distinguished from REM and NREM sleep based on manual marks made during EEG acquisition sessions and on video recordings, and considering sustained eye openings and movements. Only NREM sleep periods were retained for the following analyses. Infants with fewer than 40s of continuous and artefact-free NREM were excluded from subsequent analyses.

### EEG preprocessing

EEG data during NREM sleep were filtered using a finite impulse response (FIR) bandpass filter between 0.2 and 45 Hz. Preprocessing was then performed using the automated pipeline for infant continuous EEG (APICE) [42]. APICE combines several artifact-detection algorithms, each sensitive to specific signal features, including power spectrum, inter-channel correlation, amplitude, variance, running averages, and transient changes. These algorithms were applied iteratively to detect and correct artifacts, using adaptive thresholds that account for the high inter-individual variability observed in developing populations. After artifact detection, “bad times” were defined as periods during which 30% or more of electrodes were contaminated, while “bad channels” were identified as channels affected during more than 30% of the remaining good time data. A first correction step was applied using targeted principal component analysis to remove transient artifacts lasting less than 100 ms. This was followed by spatial interpolation using spherical splines, first locally on specific channels during remaining bad periods, and then across all “bad channels” for the entire recording. As an iterative refinement step, artifact detection was repeated to update the identification of residual bad time segments and bad channels. Bad time segments and good-time segments shorter than 8s were discarded, and any remaining artifacts were interpolated again using spherical splines. We concatenated consecutive good-time segments initially separated by bad-time intervals, marking each junction with a merge marker to ensure that this reconstructed continuity was handled appropriately in subsequent analyses. Finally, all EEG recordings were down-sampled to 250 Hz and re-referenced to the common average before further processing.

### Sleep spindles detection

Sleep spindle detection was performed on central (right: E93, E98, E103, C4 ; left: C3, E41, E42, E47), frontal (F: E4, Fz, E19), and occipital electrodes (O: E70, Oz, E83) to capture an overall view of their characteristics across the scalp. Spindle events were detected on these channels using a wavelet-based algorithm developed for adults [10,43] and adapted to the characteristics of infant sleep spindles as described below.

Candidate sleep spindles were detected by computing continuous wavelet transforms across the 11-16 Hz frequency range in 0.5 Hz steps, using a frequency B-spline wavelet (fbsp1-1-3). A dynamic thresholding algorithm based on the median absolute deviation (MAD) of the power signal was then applied: an initial hard threshold (set to 8xMAD) to isolate primary events, followed by a soft threshold (set to 4xMAD instead of 5s in adults) to precisely refine the onset and offset boundaries of each candidate. Events were officially validated as sleep spindles only if they lasted between 0.3 and 8s (instead of 2s maximum in adults), contained at least 3 oscillatory cycles, and exhibited a maximum power within the 11–16 Hz band that exceeded the power in the neighbouring 8–11 Hz and 16–19Hz frequency bands (see [10,43] for more details). The upper duration limit was chosen based on previous studies showing that sleep spindles duration is approximately 3s on average and can last up to 6s in 1-4 months old infants [9,44].Finally, detected spindles were merged if they were separated by less than 1s. If merging resulted in a duration exceeding the 8s maximum, only the longest spindle was retained and the others were discarded. Spindles that were isolated from others by more than 60s, as well as spindles overlapping a merge marker, were also discarded. Infants with fewer than 3 detected spindles were excluded from the subsequent analyses.

Several spindle features were then extracted for each infant, including inter-train interval (ITI) (s), inter-spindle interval (ISI) (s) to provide exploratory descriptions of the infraslow and mesoscale rhythms, duration (s), density (number of spindles per minute), peak frequency (Hz), amplitude (µV). Spindles were considered to belong to the same train when separated by less than 15s (based on the analyses of the section “temporal organisation of sleep spindles”, see below). ISI and ITI were calculated between spindles and spindle trains, respectively, only when no merge marker was present in between.

### Statistical analyses on sleep spindles features

The sleep spindle features extracted after detection were analysed at the cluster level by grouping adjacent electrodes. For each electrode, feature values were first averaged across all detected spindles for each infant, and then across electrodes within each cluster. Four a priori topographically defined clusters were used: a right-central (RC, E93-E98-E103-C4), a left-central (LC, C3-E41-E42-E47), an occipital (O, E70-Oz-E83) and a frontal (F, E4-Fz-E19) cluster. This cluster-based approach was chosen to reduce the impact of potential variability in EEG cap positioning between infants, which could otherwise affect the spatial comparability of the recordings. To ensure that the results were not driven by infants with marked brain abnormalities, infants with high Kidokoro scores were screened for outlying spindle-feature values. Infants with values exceeding a z-score of ≥ |2| for a given feature were excluded from the corresponding statistical analyses. For analyses of spindle features, the number of infants included thus varied across clusters of electrodes.

We aimed to determine whether sleep spindle features differed between groups (full-terms, untrained and trained preterms), and whether these group differences varied across brain regions (across the four RC, LC, O, and F clusters). After exclusion of outliers, a linear mixed-effects model (LMM) was fitted separately for each spindle feature. Group, cluster, and their interaction were included as fixed effects, with participants included as a random effect to account for the non-independence of repeated measurements across clusters within each infant. Post hoc tests were conducted using pairwise comparisons of estimated marginal means (EMMs).

### Temporal organisation of sleep spindles

We also investigated whether sleep spindles at 2M of age already exhibit the temporal clustering and rhythmic organisation previously described in adults (see [5] for a review). To do so, we adapted methods from protocols established in adult studies [43,45], focusing on the mesoscale rhythmicity, defined here as intermediate-scale temporal fluctuations in spindle activity within the 0.01-0.6 Hz frequency range. We used two complementary approaches. First, we performed a spectral power analysis in the 11-16Hz frequency band entirely independent of the previous spindle detection. Second, we used our detected spindles to generate time-frequency maps illustrating the precise interval between consecutive spindles within the same train. These analyses were performed on electrode C4. This choice is consistent with previous studies showing that sleep spindles in 2M infants are predominantly observed over frontopolar and central regions [9]. We focused on a central electrode because of its closer association with sensorimotor cortical regions, this being particularly relevant in the context of locomotor training (The results were comparable for spectral power analyses on electrodes C3, Fz and Oz, see Supplementary Information, Figure 2).

#### Spectral power analyses

For spectral power analyses, a 4-cycle Morlet wavelet centered on 11–16 Hz was applied to the entire recording to obtain the temporal envelope of spindle power. The resulting signal was then segmented into continuous NREM sleep epochs of at least 65s (vs. 30s in adults), free of merge markers (except when separated by <5 s, see Supplementary Information, Figure 3 for more details). Each epoch-specific envelope was normalised by its mean spindle power value, then convolved with a second 3-cycles (vs. 4-cycles in adults) Morlet wavelet centred on 0.01–0.6 Hz to capture mesoscale-frequency fluctuations in spindle-band activity. Epochs were averaged for each infant, weighted by duration, to obtain a mean individual low-frequency spectrum.

A one-way ANOVA with group (full-terms, untrained and trained preterms) as the between-subject factor was then performed to determine whether the mesoscale spectra differed significantly between groups. The average spectrum across infants was computed by weighting each subject according to the number of ISIs available, and infants without ISI data were excluded from this averaging procedure. Group-level average spectra were then computed separately for each group, as well as across all groups combined.

#### Time frequency analyses

Time–frequency decomposition of spindle-band power was then performed in Brainstorm [46] to further examine rhythmicity between spindles. To capture the temporal structure of spindle rhythmicity revealed by the spectral power analyses described above, segments spanning –12 to +15s around each detected spindle onset were defined. Only continuous segments free of merge markers were included. Infants were included in the subsequent analyses if at least three valid segments could be defined.

Morlet wavelets were applied to these segments to estimate time–frequency power across the 1–30 Hz frequency range, enabling the assessment of potential mesoscale rhythms. The period from −2 to −0.5s preceding spindle onset was defined as the baseline. Since all epochs were aligned to a detected spindle onset (t=0), this interval was expected to contain a low probability of spindle occurrence. Power in the resulting time–frequency maps were normalised to the baseline and then averaged for each infant. To identify significant spindle-band activity within this time window, Student’s t-tests against baseline were performed across all three groups combined and within each group separately. False discovery rate (FDR) correction was used to account for multiple comparisons.

### Spatial distribution of sleep spindles

Building on the time–frequency analyses described above, spatial topographies of spindle-band activity were assessed using time–frequency maps computed across all scalp electrodes using C4-detected spindle onsets as temporal anchors (−12 to +15 s around C4-detected spindle onsets, over 1–30 Hz). Mean power maps were extracted over the 11–16Hz frequency band for two time-windows: 0–1.5s and 1.5–3s after spindle onset. These windows were defined to capture the early and late phases of the spindle period. This total 0–3s duration aligns with the average spindle length (∼3s, though some extend up to 8s). For each window, maps were first averaged for each infant and then across infants within each group (full-terms, untrained and trained preterms), resulting in one group-level spindle-band activity map for each phase. This procedure was applied to all scalp electrodes.

To evaluate the spatial distribution of significant spindle-band activity over time, Student’s t-tests against baseline were conducted for each electrode on the average 11–16Hz power, with FDR correction for multiple comparisons. This approach yielded topographical t-maps at 0, 1, 2, and 3s for each group.

Finally, group differences in spindle-band activity were assessed using permutation testing (1000 permutations). Pairwise group comparisons were performed between the three groups at each scalp electrode for the early and late spindle windows, defined as 0–1.5s and 1.5–3s after spindle onset, respectively, averaged across the 11–16 Hz frequency range. This analysis yielded one topographical t-map for each group contrast.

## Results

A total of 50 infants had sufficient NREM sleep data for spindle detection, with a mean NREM recording duration of 286 ± 196s (range: 43 – 988s). Within this sample, mean NREM recording duration was 230 ± 133s in full-term infants (range: 56 – 511s; n = 12), 326 ± 230s in untrained preterm infants (range: 43s – 988s; n = 26), and 254 ± 159s in trained preterm infants (range: 52 – 480s; n = 12). A Kruskal-Wallis test confirmed that recording times did not differ significantly between groups (K = 1.51, p = 0.47).

### Sleep spindles features in infants

After exclusion of outliers (z score ≥ |2|), between 8 and 12 full-term infants were analysed, as well as between 17 and 26 untrained preterms, and between 11 and 12 infants trained preterms, depending on the cluster considered. Pooled across all electrode clusters and groups, the value ranges for the six features were as follows: ITI (24.35-164.65s), ISI (2.55-12.91s), duration (0.44-4.29s, see Supplementary Information, Figure 5), density (0.6-5.4 spindles/min), peak frequency (12.46-14.81Hz, with a predominance of fast spindles (13–16 Hz), see Supplementary Information, Figure 6), and amplitude (7.14-38.05µV). Mean and range of values for each feature, cluster, and group are presented in Figure 1 and detailed in the Supplementary Information, Table 3.

**Figure 1.**
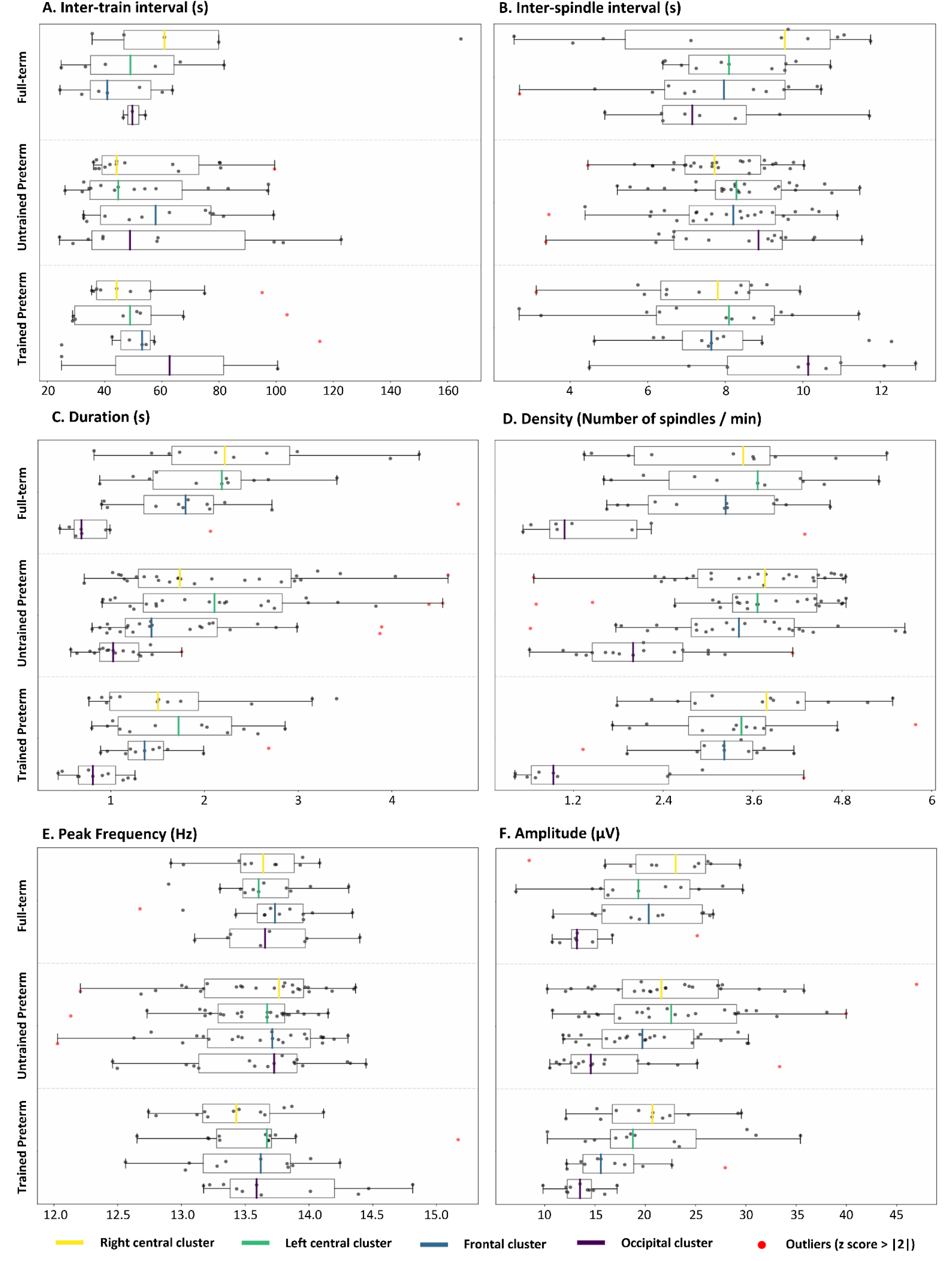
Spindles features across electrodes clusters. Quantification of sleep spindle features and identification of outliers (z-score ≥ |2|) were performed within each group (full-term, trained preterm, and untrained preterm infants) and for each electrode cluster (right-central, left-central, occipital, and frontal). Each boxplot represents the mean value per infant within each group and cluster for all analysed spindle features. **A**. Boxplot of inter-spindle-interval (ISI). **B.** Boxplot of inter-train interval (ITI). **C.** Boxplot of ratio of trains. **D.** Boxplot of density. **E.** Boxplot of amplitude. **F.** Boxplot of duration. **G.** Boxplot of peak frequency.

We first aimed to assess whether spindle features varied as a function of electrode clusters and infant groups. LMMs applied to ITI and ISI revealed no significant main effects of group or cluster, nor a significant group × cluster interaction. However, the model for spindle duration showed a significant main effect of cluster (*F* = 43.26, *p* = 2.10^-16^). The model for spindle density indicated significant main effects of group (*F* = 3.38, *p* = 0.04) and cluster (*F* = 64.36, *p* = 2.10^-16^). Finally, the model for peak frequency revealed a significant main effect of cluster (*F* = 2.96, *p* = 0.04), as well as a significant group × cluster interaction (*F* = 2.24, *p* = 0.04). For more details on the statistical results, see Supplementary Information, Table 4.

Post hoc analyses based on estimated marginal means (EMMs; see Supplementary Information, Tables 5, 6, 7, 8 for more details) confirmed a consistent effect of the occipital cluster, which differed significantly from the three other clusters for duration (*p* < 10^-4^), density (*p* < 10^-4^), and amplitude (*p* < 10^-4^). For peak frequency, the occipital cluster differed significantly only from the right-central cluster (*p* = 0.04). Across all features, the occipital cluster consistently exhibited lower values compared to the other clusters. In addition, spindle duration was significantly longer in the right-central cluster and left-central cluster than in the frontal cluster (*p* = 0.0002, and *p* = 0.004, respectively). No significant group effect on spindle density remained after FDR correction. However, a trend suggested higher spindle density in untrained preterm infants compared with full-term infants (*p* = 0.05). Finally, post hoc tests on the group × cluster interaction revealed that trained preterm infants showed significantly lower peak frequency in the occipital cluster than in the three other clusters (*p* ≤ 0.02).

### Temporal organisation of sleep spindles

#### Spectral power analyses

For these analyses, a total of 38 infants were included, comprising 9 full-term infants, 19 untrained preterm infants, and 10 trained preterm infants. One-way ANOVA with group as the between-subject factor revealed no significant group differences in peak frequency of the mesoscale power spectrum: *F*(2,35) = 1.01, *p* = 0.38 (see Supplementary Information, Figure 4). Accordingly, a mean mesoscale spectrum was computed across all 38 infants. This group-averaged spectrum reached its maximum at 0.08 Hz, corresponding to a period of 12.5s (Figure 2 - A). Across infants, individual peak frequencies ranged from 0.02 to 0.14 Hz.

**Figure 2.**
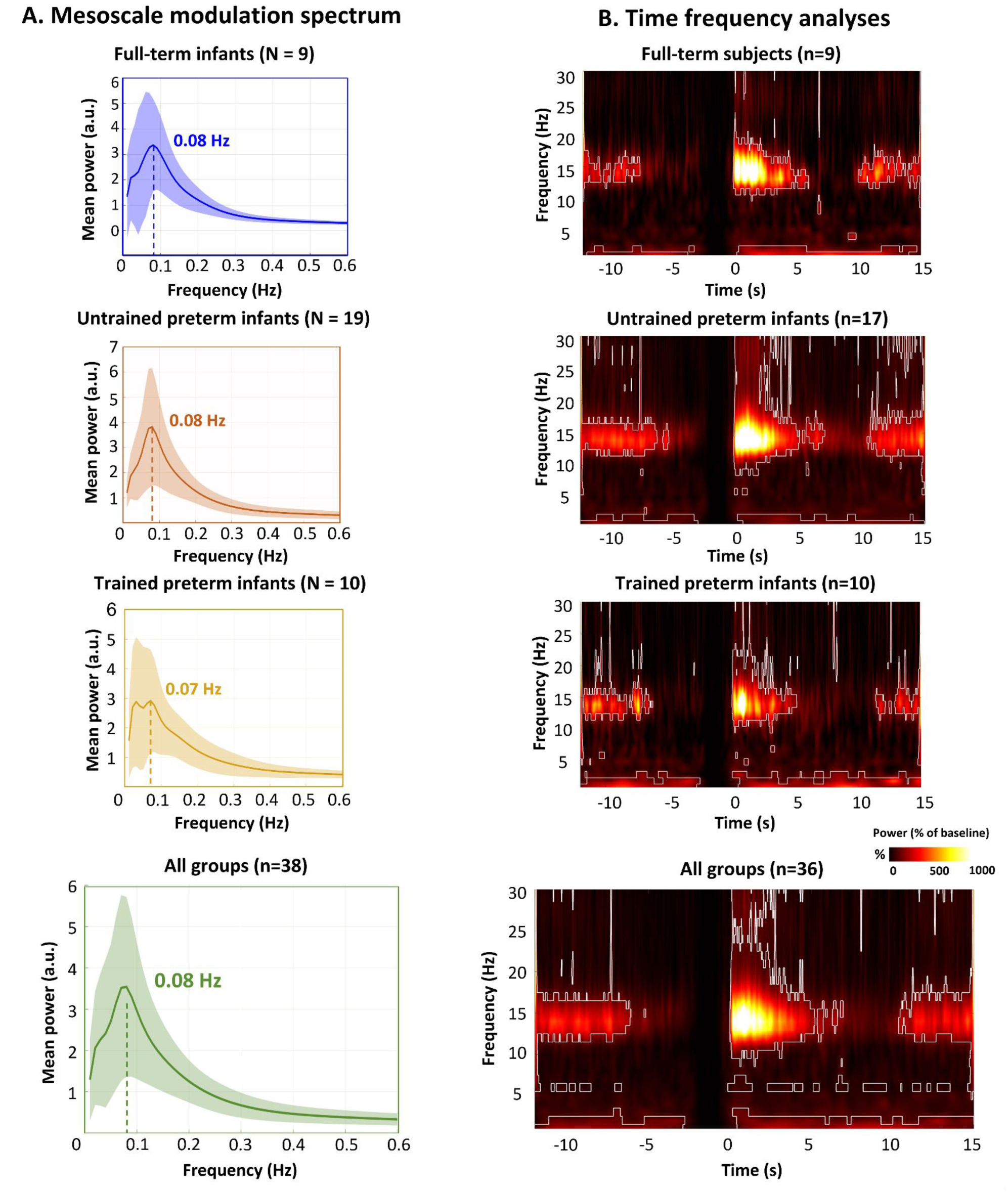
Spindle temporal organisation (C4) **A.** Grand-average spectral profile recorded at electrode C4, computed across all three groups and separately for each group. The grand-average spectrum was weighted by the number of inter-spindle intervals contributed by each participant. It illustrates fluctuations in spindle-band power (11–16 Hz) over a low-frequency range from 0.01 to 0.6Hz, reflecting mesoscale modulation. A prominent peak at 0.08 Hz indicates a periodic structure in spindle-band activity at this timescale. The shaded light-green area around the mean curve represents the standard deviation across participants. **B.** Grand average TF maps zoomed in on the −12 to +15 s window around spindle onsets at C4, illustrating spindle mesoscale modulation. The colour bar reflects spectral power relative to baseline (%). Significant clusters (p < 10⁻³⁰) identified using FDR-corrected t-tests are outlined in white. TF maps were computed for full-term, trained preterm, and untrained preterm infants, as well as for all groups combined, as no significant group differences were observed.

#### Time frequency analyses

The time-frequency decomposition of spindle band-power revealed results consistent with those obtained from the mesoscale power spectrum analysis. A total of 36 infants were included in these analyses, comprising 9 full-term infants, 17 untrained and 10 trained preterm. As shown in Figure 2 - B, spindle-band activity exhibited a rhythmic organisation that differed significantly from baseline (*p* < 10⁻³⁰), with spindles occurring on average approximately every 12 seconds.

### Spatial distribution of sleep spindles

The spatial topographies of spindle-band activity are presented in Figure 3. The mean power maps normalised relative to baseline (Figure 3 – A), together with t-maps showing spindle activity significantly different from baseline (Figure 3 – B), revealed that spindle activity was mainly localised over central and frontal regions across the three groups.

**Figure 3.**
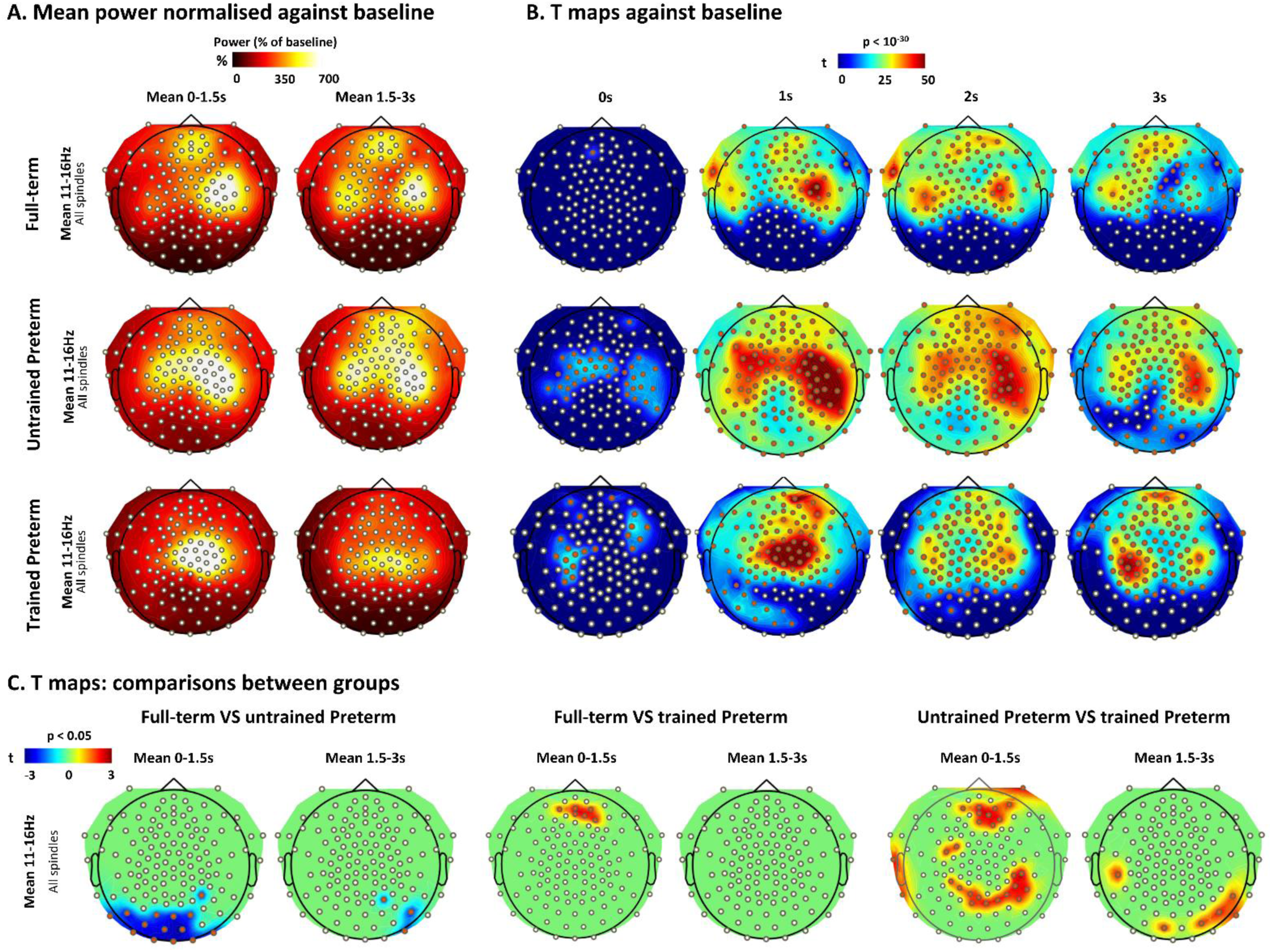
Spatial topography of spindle-band activity. **A.** Spatial topography maps of spindle-band power normalised relative to baseline and averaged over two time windows following spindle onset detection at C4: 0–1.5s and 1.5–3s. **B.** FDR-corrected t-tests against baseline performed on the averaged spindle-frequency band at specific time points (0, 1, 2, and 3 s). **C.** Permutation test results obtained from all pairwise comparisons between the three groups. All channels highlighted in red correspond to channels exhibiting a significant effect.

#### Effect of prematurity: Untrained preterm infants versus full-term infants

Figure 3 – A highlights a more diffuse spindle activity pattern in untrained preterm infants compared with the more localised organisation observed in full-term infants, which was characterised by distinct clusters in the left central, right central, and frontal regions. This reduced spatial specificity of activation in untrained preterm infants is also reflected in Figure 3 – B, which shows spindle-band activity significantly different from baseline (p < 10^-30^) across all channels, whereas activity in full-term infants appeared more spatially restricted. This pattern of activation was consistently observed across all analysed time periods. Similarly, group comparisons shown in Figure 3 – C revealed significantly higher spindle activity in occipital regions in untrained preterm infants compared with full-term infants (p < 0.05), particularly during the early spindle window from 0 to 1.5 s after spindle onset.

#### Effect of training: Trained preterm infants versus full-term infants and untrained preterm infants

Compared with untrained preterm infants, trained preterms displayed a more localised distribution of spindle activity. Rather than exhibiting a diffuse scalp-wide pattern, spindle-band activity in trained preterm infants displayed a more spatially defined organisation. However, this topography lacked the sharply defined spatial clustering of full-term infants. As shown in Figure 3 – A, activity was primarily concentrated over central regions, while Figure 3 – B revealed the progressive emergence of a frontal cluster significantly different from baseline (*p* < 10⁻³⁰). Accordingly, group comparisons shown in Figure 3 – C indicated significantly lower spindle activity in trained than in untrained preterm infants over frontal and parietal regions during the early spindle window, from 0 to 1.5 s after spindle onset. During the later window, from 1.5 to 3 s after spindle onset, trained preterm infants also showed lower occipital activity than untrained preterm infants. Finally, although a frontal activation cluster was observed in trained preterm infants (Figure 3 – B), their frontal spindle activity remained significantly lower than that of full-term infants.

## Discussion

This study provides an original investigation of sleep spindles at their emergence period, drawing on a dataset of three distinct groups of infants assessed during sleep at 2 months of (corrected) age: full-term infants, very preterm infants, and very preterm infants receiving early locomotor intervention. This three-group comparison allowed us to explore both typical and atypical development of spindle characteristics, while assessing the effect of an early locomotor intervention on spindle development and organisation.

Two main findings emerged. First, we highlighted a rhythmic pattern of sleep spindle occurrence across all three groups of infants, operating at a timescale approximately three times slower than that observed in adults. Second, sleep spindle spatial distribution differed significantly across groups. Untrained preterm infants displayed a more diffuse scalp distribution compared to the other two groups, while trained preterm infants showed a more focal distribution of spindle activity, predominantly over central regions, but with reduced frontal activity relative to both full-term and untrained preterm infants.

### Early sleep spindles features in full-term and preterm infants

First, spatial topography maps revealed particularly pronounced sleep spindle activity over central and frontal regions across the three groups. This result was consistent with our statistical analyses of sleep spindle characteristics, which revealed consistently lower duration, density, and amplitude in the occipital cluster compared with the other clusters. These findings are in line with previous studies conducted on full-term infants at a similar age [9,47] or a few months later ([13], see [15] for a review). With a central spindle density (median = 3.6 spindles/min) similar to that of adults (3.07 spindles/min ; [48]), these findings suggest that the fronto-central predominance of sleep spindles emerges early in development and is already observed in very preterm infants. This spatial consistency may reflect the early regional organisation of the thalamo-cortical circuits involved in spindle generation and propagation.

We also highlighted the presence of a mesoscale rhythm with a period of ∼12.5 s (peak frequency of approximately 0.08 Hz) across the three groups, approximately three times slower than the ∼4-s recurrence period reported in adults ([43], see [5] for a review). Importantly, this temporal organisation was observed not only in the spectral analyses but also in the time-frequency maps aligned to spindle onsets. These converging analyses therefore suggest that a structured temporal organisation of spindle activity is already present at 2M, although its characteristic timescale differs markedly from that observed in the mature brain. One possible explanation for this slower recurrence is the progressive maturation of thalamo-cortical networks generating sleep spindles. This slower mesoscale rhythm may reflect the immaturity of thalamo-cortical circuits, including reduced conduction and connectivity due to incomplete myelination, immaturity of microstructure and other mechanisms regulating spindle initiation and termination within the TRN (thalamic reticular nucleus) – TC (thalamocortical neurons) loop (see [15,49,50] for reviews on the neurophysiological mechanisms underlying sleep spindles). However, this interpretation remains speculative, since scalp EEG does not allow us to directly identify the cellular mechanisms underlying spindle timing and organisation.

To our knowledge, only one previous exploratory study had specifically investigated spindle rhythmicity in young infants. Using MEG in seven full-term infants aged between 46 and 63 weeks, [11] reported a clock-like recurrence of sleep spindles at a frequency of approximately 0.1 Hz. This recurrence rate is slightly faster than the 0.08-Hz rhythm identified in our study, which is consistent with the fact that some of the infants in that study were older than those included in ours. The broader and partly older age range of their sample may contribute to this difference and raises the possibility that spindle recurrence progressively accelerates during early development, a hypothesis that will require longitudinal investigation.

In another study, [9] also estimated the average minimal refractory period between sleep spindles detected at electrodes C3 and C4, reporting an ISI of approximately 5s. However, their ISI was intended to approximate the minimal recovery time required for thalamo-cortical circuits to generate a new spindle, based on the 10% shortest intervals, which may partly explain the shorter refractory period they reported compared to ours. In addition, their approach differed from ours, as they defined the ISI from spindle offset to the subsequent spindle onset, whereas we measured ISI between spindle onsets and additionally relied on robust spectral and time–frequency analyses to characterise a mesoscale rhythm complementing ISI-based metrics. These methodological differences likely contribute to the shorter value reported by Kwon and colleagues. Importantly, in our study, the mean ISI derived from detected spindle events (∼8 s) was also shorter than the ∼12.5-s periodicity identified by spectral and time–frequency analyses. This distinction suggests that event-to-event intervals and rhythmic modulation of spindle-band activity capture related but non-equivalent aspects of spindle temporal organisation, highlighting the complementarity of event-based, spectral, and time-frequency approaches for characterising spindle temporal dynamics. Although spectral and time-frequency analyses could not be performed to identify the infraslow rhythm, the ITI identified based on spindle detection (range: 43.60-77.57s) was comparable to that reported in adults (∼50s, e.g. [43,45], see [5] for a review), or even longer depending on the groups and clusters considered.

This slower rhythm observed in infants compared with adults is also consistent with relatively longer spindle durations detected, ranging from approximately 0.3 to 8 s, compared with the ∼0.3-2 s range generally reported in adults (e.g. [10,43]), with both findings pointing toward a possible reduction in the efficiency of the TRN–TC loop. Longer spindle durations during early infancy have also been described previously [9,44,51], although the reported upper limits varied across studies. Together, these observations suggest that both the duration and temporal recurrence of spindle activity undergo substantial maturation during the first months of life.

The coupling between slow oscillations (0.5–1.25 Hz) and sleep spindles is also a developmental marker known to reflect coordinated activity within thalamo-cortical and hippocampal networks, and has been proposed to play a crucial role in effective memory consolidation in adults ([43,52], see [5,49] for reviews). Hence, to further test the hypothesis of still immature thalamo-cortical networks in 2m-old infants, we further performed an exploratory analysis to investigate the coupling between slow oscillations and sleep spindles after pooling the three infant groups (see Supplementary Information, Figure 7). We did not identify any significant slow oscillation–spindle coupling in our dataset, which might be compatible with still immature coordination between slow oscillations and sleep spindles at 2 months of age. Nevertheless, this exploratory analysis should not be interpreted as direct evidence of an inefficient thalamo-cortical network since it was constrained by the limited amount of NREM sleep periods. Longer sleep recordings and longitudinal measurements will be necessary to establish when robust slow oscillation-spindle coupling emerges during infancy and how it relates to the maturation of thalamo-cortical networks.

### Impact of prematurity on sleep spindle features

At the cluster level, only a trend toward higher spindle density was observed in extreme and very preterm infants (untrained) compared with full-term infants. No other effects of prematurity were found on spindle characteristics extracted using our detection algorithm, including ITI, ISI, and spindle duration. Overall, these findings suggest that the characteristics of individual spindle events are broadly comparable between full-term and very preterm infants at 2 months of corrected age. Interestingly, however, prematurity was associated with clear differences in the spatial distribution of spindle-band activity. Untrained preterm infants displayed a more diffuse scalp-wide pattern than full-term infants, whose activity was organised into more clearly defined frontal and bilateral central clusters. In addition, preterm infants showed significantly greater occipital spindle-band activity during the early window following spindle onset, consistent with a more widespread spatial distribution of spindle activity across the scalp. No corresponding group differences were detected in terms of the mesoscale temporal organisation of spindles. Taken together, these results suggest that prematurity-related differences are more apparent in the spatial distribution of spindle-related activity than in its temporal organisation. [53] demonstrated that premature birth impairs structural brain network efficiency by selectively reducing short-range fiber density while preserving long-range rich-club connections. Crucially, by enabling local cortical GABAergic inhibition of thalamic excitation, short-range connections constrain spindle propagation (see [15,49] for reviews). A deficit in short-range connectivity may therefore contribute to the overly diffuse spindle activity observed in preterm infants.

The increased spindle-band activity across the scalp, particularly in the occipital region, may reflect accelerated developmental processes in the visual cortex of preterm infants, potentially driven by earlier exposure to the extra-uterine environment. Consistent with this hypothesis, studies investigating different neurophysiological markers of maturation in this region have converged on similar conclusions in infants born preterm. [54], for example, reported evidence of faster visual evoked potentials in very preterm infants, while [55] observed more advanced functional connectivity in sensory regions—including occipital areas— in ex-utero preterm infants compared to in-utero foetuses of equivalent gestational age. [56] similarly reported flatter aperiodic EEG slopes—particularly over occipital regions— an index often interpreted as reflecting a more mature neural profile. However, this apparent acceleration of visual system maturation with prematurity should be interpreted with caution, as its functional significance remains uncertain. Earlier sensory exposure could induce adaptive changes without necessarily promoting more favorable neurodevelopment. Indeed, [54] emphasised that accelerated maturation of visual evoked responses may reflect adaptation to earlier extra-uterine visual stimulation, and has in some cases been associated with less favourable long-term cognitive outcomes [57]. Therefore, the increased occipital activity observed in preterm infants in our study may reflect altered neurodevelopmental organisation rather than advanced maturation.

A complementary interpretation is that the more diffuse sleep spindle activity observed in preterm infants may reflect reduced regional specialisation of the networks involved in early sensorimotor processing and potentially in subsequent memory consolidation. This interpretation is consistent with previous fMRI studies in adults showing that early stages of motor learning and lower levels of expertise are associated with broader and less efficient patterns of brain activation. For instance, compared with trained sequences of finger movements, untrained movement sequences evoked broader activation across motor and parietal regions, whereas training led to more focal and sequence-specific neural representations [58]. Similarly, two fMRI studies investigating both motor execution and motor imagery reached the same conclusion. [59] reported that individuals with poorer motor imagery ability exhibited broader bilateral activation suggesting greater neural effort to perform the task. Likewise, [60] showed that expert violinists displayed highly focal activation patterns during both motor execution and motor imagery, whereas amateur musicians recruited a much more widespread network. Although these adult findings cannot be directly extrapolated to the developing infant brain, they provide a useful conceptual framework in which motor expertise is characterised by greater neural efficiency, reflected in more selective recruitment of task-relevant regions. On the reverse, less-skilled adult individuals rely on more diffuse brain activation patterns that likely reflect less automated and more resource-demanding processing. Given that the cortical distribution of sleep spindles preferentially involves regions engaged during prior learning ([4,61,62], see [5] for a review), the diffuse spindle-band distribution observed in preterm infants may therefore reflect a lower degree of regional specialisation within the developing networks contributing to sensorimotor processing and sleep-dependent plasticity. Although high-density EEG recordings provide reasonable topographical resolution and can help to infer distribution patterns, future multimodal studies combining EEG with structural or functional MRI would be valuable to further refine the anatomical specificity of these effects and confirm their cortical localisation.

### Early locomotor training and sleep spindle modulation in preterm infants

No significant differences in the temporal organisation of sleep spindles were observed between untrained preterm infants and those who underwent early training on the Crawliskate at home for 2 months. This suggests that the temporal rhythmicity of spindle activity may be independent of prior sensorimotor experience. This interpretation is consistent with the findings of [43], who reported no differences in spindle rhythmicity between adults who underwent arm immobilisation during the day preceding the recorded sleep period and those who did not.

However, the trained group showed a markedly different spatial organisation. Rather than showing the diffuse scalp-wide pattern observed in the untrained group, trained preterm infants exhibited a more focal distribution of spindle activity, predominantly over central regions, resembling the spatially localised organisation observed in full-term infants. However, the spatial pattern observed in the trained group remained distinct from that of full-term infants, with a central activity that remained slightly broader compared with the two distinct bilateral central clusters observed in full-term infants. In addition, frontal sleep spindle activity was significantly lower in the trained group than in both full-term and untrained preterm groups. Interestingly, our findings suggest that early locomotor experience in very preterm infants may be associated with changes in the spatial organisation of sleep spindle activity, with a more central distribution that may reflect greater regional specialisation over sensorimotor cortical regions. This central predominance is particularly noteworthy given the sensorimotor nature of the Crawliskate intervention. As sleep spindles emerge from developing thalamo-cortical networks, their spatial expression may be shaped by early sensorimotor experiences. During early infancy, spontaneous movements and sensorimotor interactions contribute to the formation of sensorimotor cortical maps. In this context, repeated locomotor experience, such as the one with the Crawliskate, may influence the developing networks engaged during sensorimotor activity and their subsequent expression during sleep, potentially leading to a more focal distribution of spindles. The lower occipital spindle-band activity observed in trained compared with untrained preterm infants is also compatible with a shift toward a more spatially differentiated pattern. However, the present data do not allow us to determine whether early sensorimotor training induced a shift in spindle activity from occipital toward central regions, or whether the observed pattern reflects a more specialised spatial organisation. Hence, despite earlier extra-uterine sensory exposure in both preterm groups, including the visual modality, greater sensorimotor experience may contribute to a more selective recruitment of regions involved in sensorimotor processing, accompanied by reduced occipital sleep spindle activity. Future longitudinal within-subject studies could test whether this spatial reorganisation reflects experience-dependent plasticity by assessing spindle organisation before and after locomotor training at an age when sleep spindles can be reliably identified. However, initiating training at this later stage may capture a reduced window of developmental plasticity.

These findings are further supported by the study of [32], who reported improvements in global motor performance in preterm infants following Crawliskate training at 2, 6, 9 and 12M. Better motor outcomes following locomotor training may reflect more advanced development of sensorimotor networks. Given the proposed role of sleep spindles as markers of motor learning, these findings further suggest that locomotor training may contribute to optimising the spatial organisation of sleep spindles. Moreover, the reduced frontal sleep spindle activity observed in trained preterm infants compared with both full-term and untrained preterm infants may also reflect a spatial distribution of spindle activity that more closely resembles a later developmental stage: in a large developmental study spanning infancy to adolescence, the most pronounced decrease in spindle rate from 4 to 10 months of age was observed over frontal regions [9]. Future longitudinal studies should investigate whether these early changes persist over time and examine their relationship with the acquisition of motor milestones during infancy to provide further insight into the functional significance of these developmental changes.

### Limitations

Despite its major findings, several limitations of this study should be acknowledged. First, although this represents a valuable longitudinal dataset including an intervention in a subgroup of preterm infants, this spindle analysis primarily compared groups at a single developmental time point. The sample sizes might seem relatively small and unbalanced across groups, mainly due to the difficulty in recording NREM sleep in infants in a research setting, and to train infants at home with parents. Replication in larger samples with longitudinal EEG recordings at several developmental stages will therefore be important, ideally using longer sleep recordings, such as full-night recordings, to improve the robustness of the findings. Second, although no significant differences in GA were observed between preterm groups, trained infants were on average slightly older than untrained infants at EEG recording. Given the rapid maturation occurring during this developmental period, this remains a limitation, though supplementary analyses in age-matched subgroups yielded the same main effects as the whole-group analysis. Third, the preterm population was clinically heterogeneous, including the brain abnormalities with a trend toward slightly higher severity (i.e. higher Kidokoro scores) in the trained preterm group. Although this heterogeneity improves the ecological validity of the sample by better reflecting the clinical population, the current sample was too small to disentangle the respective contributions of prematurity versus brain abnormalities or other maturational delays. However, additional checks using a subgroup of untrained infants with homogeneous brain abnormalities as trained infants showed no effect on our main findings. Finally, the limited duration of recordings and short NREM periods in most infants constrained the ability to reliably identify the slower, infraslow organisation of spindle trains. Therefore, although their rhythmic organisation is consistent with spindle trains described in adults, it remains unclear whether consecutive spindles already form distinct trains or reflect the progressive emergence of this organisation. The inability to identify the infraslow rhythm characteristic of spindle trains in this study prevents us from distinguishing between these possibilities. We observed mesoscale oscillatory activity, but no group differences emerged at this level. However, more subtle differences in infraslow temporal organisation cannot be excluded.

## Conclusion

Using a unique cohort of infants born very preterm or at term, we characterised the early temporal and spatial organisation of sleep spindles and examined its sensitivity to prematurity and early locomotor training. No effect of preterm birth was found on the temporal organisation of sleep spindles but an effect was observed on their spatial organisation. Preterm infants exhibit a more diffuse scalp wide activity in comparison to full-terms showing a more clearly organised activity in frontal and bilateral central clusters. However, following locomotor training, preterm infants exhibited a more focal distribution—although less clearly clustered than that observed in full-term infants— predominantly over central regions. With this study, we identified a potential neural marker that may provide new insights into the mechanisms underlying sensorimotor development and help improve the characterisation and understanding of sensorimotor developmental delays in preterm infants. Furthermore, we provide preliminary evidence for the potential neurodevelopmental benefits of a locomotor intervention. As an intervention that can be implemented from a very young age, this may offer a unique opportunity to provide sensorimotor experience during a critical period of development, when early intervention may be particularly beneficial for shaping developmental trajectories in infants at risk for later neurodevelopmental difficulties.

## Acknowledgments

The authors thank the French National Agency for Research (ANR grant ANR-20-CE17-0014-03-PremaLocom), the Fondation Médisite (under the aegis of the Fondation de France, grant FdF-18-00092867), the IdEx Université de Paris (ANR-18-IDEX-0001), the French government as part of the France 2030 programme (grant ANR-23-IAIIU-0010, IHU Robert-Debré du Cerveau de l’Enfant) for their support. We gratefully acknowledge the clinical team at Robert-Debré Hospital and at NeuroSpin/UNIACT whose collaboration enabled this study to be carried out in clinical settings. Last but not least, we would like to sincerely thank all the infants and parents who participated in this study.

