## Supplementary Information for "Characterising sleep spindle emergence in infancy: Effects of prematurity and early locomotor intervention"

#### Locomotor training

**Supplementary information. Table 1. Individual subject profiles and group average over the 8-week training period**

| Subjects | Average number of sessions over 8 weeks | Average training time over 8 weeks | Average distance covered over 8 weeks | Average session speed over 8 weeks |
|---|---|---|---|---|
| **mean ± sd (range)** | 39,8 ± 10,5<br>(19 – 53) | 358,0 ± 107,8<br>(182,3 - 493,9) | 211,4 ± 90,3<br>(77,4 - 334,4) | 0,6 ± 0,2<br>(0,4 -0,8) |

#### Homogeneity between preterm infant groups

**Supplementary information. Table 2. Characteristics of preterm infants included in the time–frequency analyses (with an untrained subgroup, matched to the trained group).**

| **Factors** | **Untrained preterm group n(%)** | | | | **Trained preterm group (Crawliskate) n(%)** | | | |
|---|---|---|---|---|---|---|---|---|
| **N subjects (initial)**<br>**(Matched subgroup)** | 17<br>10 | | | | 10 | | | |
| **GA at birth** | G1 | G2 | G3 | | G1 | G2 | G3 | |
| **Mean (SD)** | 6 (35.3 %)<br>3 (30.0%) | 6 (35.3 %)<br>3 (30.0%) | 5 (29.4 %)<br>4 (40.0%) | | 3 (30.0 %) | 3 (30.0 %) | 4 (40.0 %) | |
| **Kidokoro score** | Normal | Mild | Moderate | Severe | Normal | Mild | Moderate | Severe |
| | 11 (64.7 %)<br>5 (50.0%) | 5 (29.4 %)<br>4 (40.0%) | 0 (0.0 %)<br>0 (0.0%) | 1(5.9 %)<br>1 (10.0%) | 5(50.0 %) | 3(30.0 %) | 1(10.0 %) | 1(10.0 %) |
| **Brain lesions** | SEH | IVH2 | IVH3 | IVH4 | SEH | IVH2 | IVH3 | IVH4 |
| | 1 (5.9 %)<br>1 (10.0%) | 4 (23.5 %)<br>2 (20.0%) | 3 (17.6 %)<br>2 (20.0%) | 0 (0.0 %)<br>0 (0.0%) | 2 (20.0 %) | 1 (10.0 %) | 1 (10.0 %) | 0 (0.0 %) |

We conducted supplementary analyses to ensure our findings were not confounded by slight heterogeneity in age and brain abnormalities among the preterm infants. Out of the 17 untrained subjects initially included in the time-frequency analyses of sleep spindle spatial and temporal organisation, we selected a size-matched subgroup of 10 infants. These infants were specifically chosen to match the trained group's clinical characteristics, including brain lesions, Kidokoro scores, and age. The optimal matched combination is presented in blue in Supplementary Information. Table 2.

The same analyses as those described in the "Spatial distribution of sleep spindles" section of the main results were subsequently performed on this subgroup of untrained infants to obtain mean power maps across two time windows: 0–1.5 s and 1.5–3 s following spindle onset. Supplementary Information Figure 1 highlights a more diffuse spindle activity pattern in untrained preterm infants replicating main group findings (see main Results, Figure 3). To ensure that the spatial organisation of the 10 untrained infants most closely matching the trained infants did not significantly differ from that of the entire untrained group, a permutation test (n=1000) was conducted comparing the mean power maps of the subgroup versus those of the full group. No significant differences were found (Supplementary Information. Figure 1). We also compared the spatial distribution of sleep spindles between the subgroup of untrained infants and the group of trained infants to determine whether the same pattern of results was observed as when considering the entire group of untrained infants, using permutation testing (n = 1000). Consistent with the main results obtained for the whole group of untrained infants, we found significantly lower spindle activity in frontal and parietal regions in the trained group during the early spindle window (0-1.5s). During the later window (1.5-3s), trained infants also showed lower occipital activity than untrained infants (Supplementary Information. Figure 1).

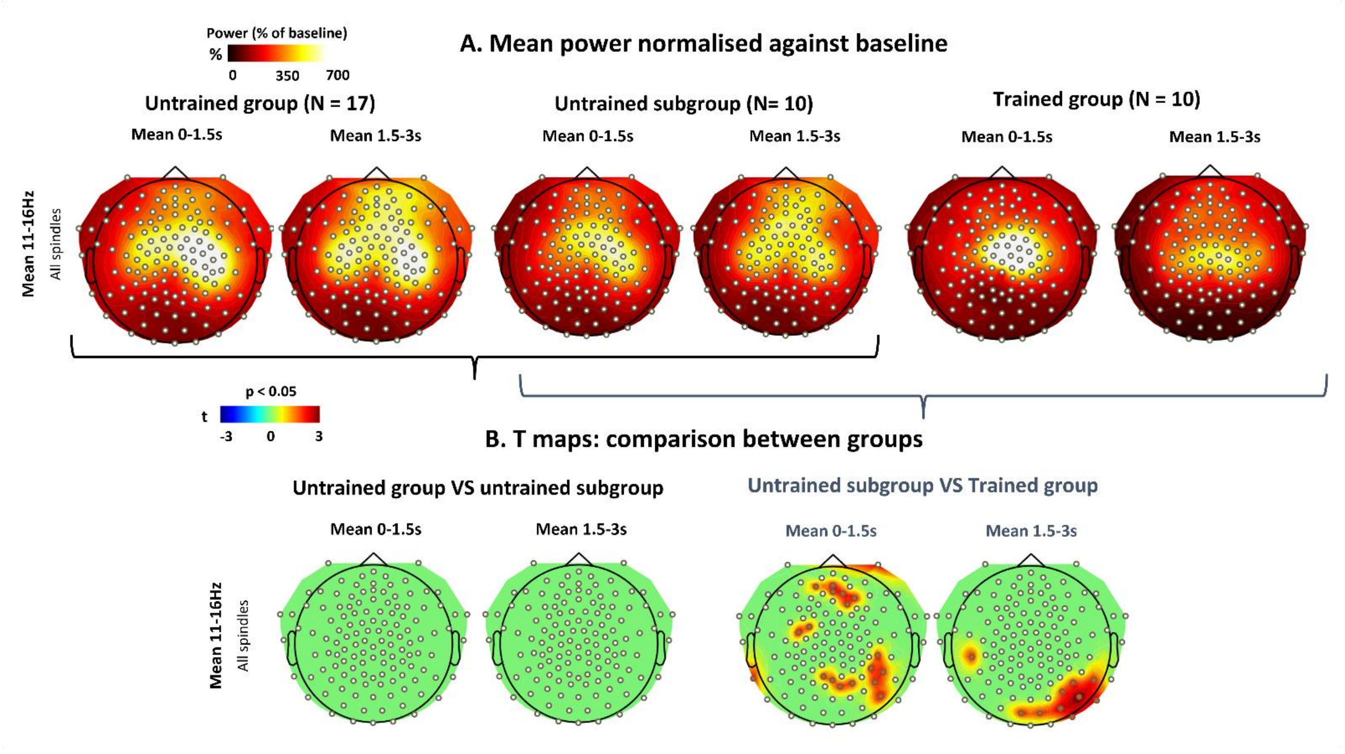


**Supplementary Information. Figure 1. Spatial distribution of sleep spindles between preterm groups: Control analysis using the untrained subgroup matched to trained infants**

**A.** Spatial topography maps of spindle-band power normalised relative to baseline and averaged over two time windows following spindle onset detection at C4: 0–1.5s and 1.5–3s. **B.** Permutation test results obtained from comparisons between all untrained infants (N = 17) and the untrained subgroup (N = 10) ; and between the untrained subgroup (N = 10) and the trained group (N = 10).

All channels highlighted in red correspond to channels exhibiting a significant effect.

### Temporal organisation of sleep spindles: Spectral power analysis

#### Mesoscale modulation spectrum across electrodes

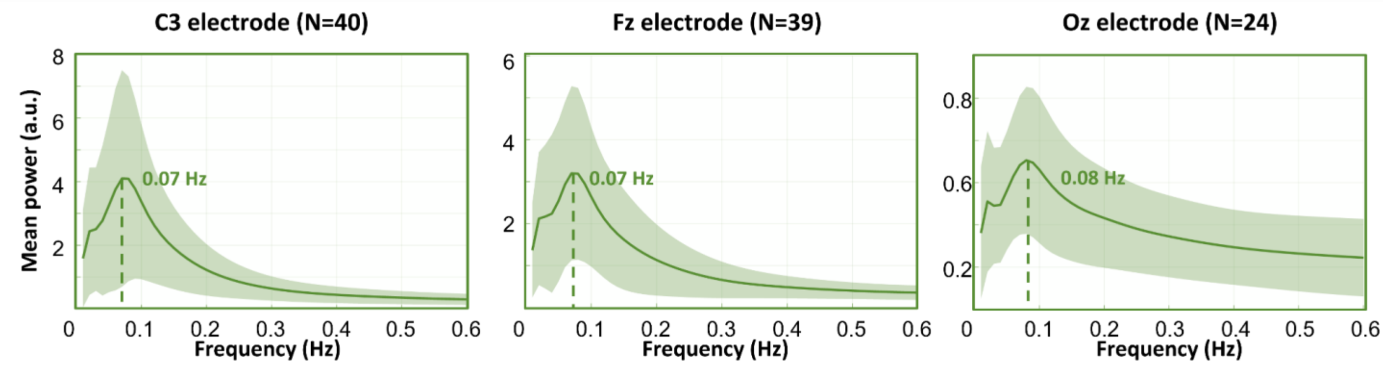


**Supplementary information. Figure 2. Mesoscale modulation spectrum on C3, Fz and Oz electrodes (all groups)**

Grand-average spectral profile recorded at electrodes C3, Fz, Oz, computed across all three groups. The grand-average spectrum was weighted by the number of inter-spindle intervals contributed by each participant. It illustrates fluctuations in spindle-band power (11–16 Hz) over a low-frequency range from 0.01 to 0.6 Hz, reflecting mesoscale modulation. A prominent peak at 0.08 Hz indicates a periodic structure in spindle-band activity at this timescale. The shaded light-green area around the mean curve represents the standard deviation across participants.

#### Validation of zero-padding for gaps < 5s between concatenated good-time segments

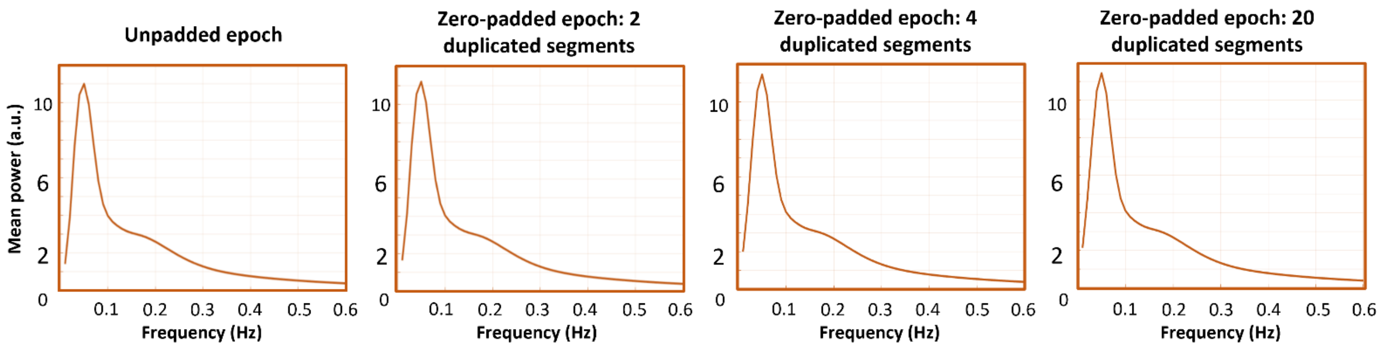


**Supplementary information. Figure 3.** Effect of zero-padding (<5s gaps) on the mesoscale power spectrum

To ensure that at least two oscillation cycles could be observed for each infant, only infants with at least 65s of continuous NREM sleep free of merge markers were included in these analyses. To maximise the use of available data, an exception to this criterion was made when two good-time segments were originally separated by less than 5s. In these cases, the segments were concatenated by filling the gap with zeros corresponding to its duration, thereby preserving a continuous representation. This 5s threshold was considered appropriate given the low likelihood of spindle occurrence within such a short interval in infants.

To ensure that this zero-padding procedure did not introduce spectral artifacts, we performed the following validation test. An artifact-free 130s epoch was selected from a subject's recording, for which no spindle had been identified within the last 5s (125-130s). This epoch was truncated to its first 125s and was then duplicated. The two copies were concatenated with a 5s zero-vector inserted between them. We applied the same procedure using four duplicated segments, and twenty segments each separated by a 5s vector of 0. The same approach was then extended to four and to twenty duplicated segments, each pair separated by a 5s zero-vector, resulting in three reconstituted test epochs. The same procedure described in the main Method was applied to each of these reconstituted epochs to obtain their mesoscale spectrum. The results were compared with the spectrum computed on the original, unpadded epoch. No distortion of the spectral shape was observed at the junctions between the real signal and the zero-padded segments: peak frequency was unaffected, with only a slight variance in peak power. These results confirm that inserting up to 5s of zeros between concatenated segments does not bias the spectral analysis.

<u>Mesoscale modulation spectrum across groups</u>

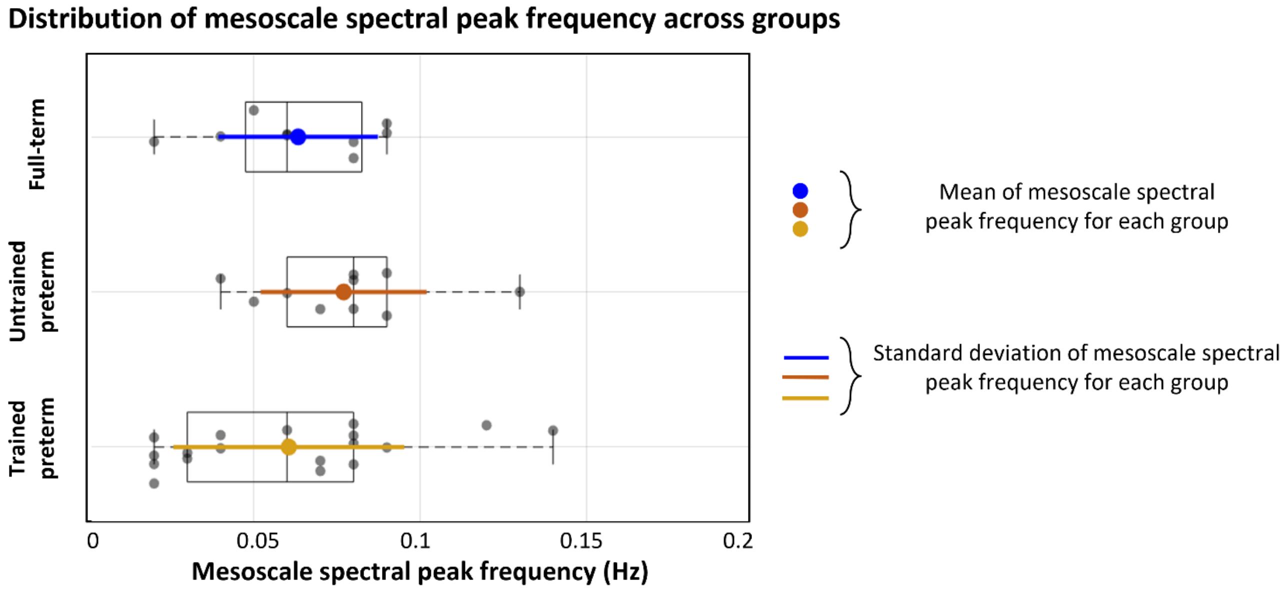


**Supplementary information. Figure 4.** Distribution of mesoscale spectral peak frequency across groups
Distribution of the mesoscale spectral peak frequency in the full-term, untrained preterm, and trained preterm groups. Peak frequencies were obtained from individual non-ISI-weighted spectra and were used for the ANOVA testing group differences in mesoscale modulation. No significant group effect was detected.

#### Sleep spindle features

**Supplementary Information. Table 3. Sleep spindles features per group.**

For each spindle feature, subjects with fewer than 3 spindles were excluded from the analysis. Subjects with a *z*-score ≥ |2| were also excluded.

| Spindle features: mean ± sd (range) | Clusters | Full-term infants | Trained preterm infants | Untrained preterm infants |
|---|---|---|---|---|
| ITI | RC | 77.57 ± 51.42 (35.62–164.65) N = 5 | 46.43 ± 13.58 (35.44–74.95) N = 8 | 52.58 ± 17.62 (36.10–80.60) N = 14 |
| | LC | 50.71 ± 21.67 (24.79–81.79) N = 6 | 43.60 ± 14.98 (28.76–67.56) N = 7 | 53.57 ± 23.87 (26.12–97.29) N = 15 |
| | F | 44.46 ± 14.67 (24.35–63.71) N = 7 | 46.86 ± 11.92 (24.92–57.40) N = 6 | 59.52 ± 22.57 (32.61–99.11) N = 12 |
| | O | 50.14 ± 3.89 (46.50–54.25) N = 3 | 62.73 ± 53.47 (24.92–100.53) N = 2 | 60.76 ± 35.04 (24.19–122.76) N = 10 |
| ISI | RC | 8.15 ± 3.28 (2.55–11.75) N = 10 | 7.72 ± 1.40 (5.75–9.93) N = 11 | 7.92 ± 1.23 (5.66–10.03) N = 25 |
| | LC | 8.35 ± 1.50 (6.39–10.71) N = 11 | 7.48 ± 2.61 (2.68–11.45) N = 12 | 8.36 ± 1.59 (5.22–11.47) N = 24 |
| | F | 8.11 ± 1.92 (4.64–10.47) N = 11 | 8.04 ± 2.26 (4.62–12.29) N = 11 | 8.15 ± 1.61 (4.38–10.89) N = 23 |
| | O | 7.67 ± 2.12 (4.89–11.72) N = 8 | 9.58 ± 2.63 (4.49–12.91) N = 9 | 8.51 ± 1.67 (6.21–11.52) N = 15 |
| Duration | RC | 2.40 ± 1.11 (0.82–4.29) N = 10 | 1.68 ± 0.89 (0.76–3.41) N = 12 | 2.01 ± 0.90 (0.71–4.04) N = 25 |
| | LC | 2.02 ± 0.73 (0.88–3.41) N = 11 | 1.72 ± 0.71 (0.80–2.86) N = 12 | 2.01 ± 0.87 (0.91–3.83) N = 22 |
| | F | 1.69 ± 0.58 (0.90–2.72) N = 11 | 1.34 ± 0.32 (0.89–1.99) N = 10 | 1.61 ± 0.64 (0.80–2.99) N = 22 |
| | O | 0.71 ± 0.20 (0.45–0.99) N = 7 | 0.84 ± 0.28 (0.44–1.26) N = 11 | 1.03 ± 0.26 (0.57–1.44) N = 16 |
| Density | RC | 3.00 ± 1.20 (1.20–5.40) N = 10 | 3.60 ± 1.20 (1.80–5.40) N = 12 | 3.60 ± 0.60 (2.40–4.80) N = 25 |
| | LC | 3.60 ± 1.20 (1.80–5.40) N = 11 | 3.00 ± 1.20 (1.80–4.80) N = 11 | 4.20 ± 0.60 (2.40–4.80) N = 22 |
| | F | 3.00 ± 1.20 (1.80–4.80) N = 12 | 3.00 ± 0.60 (1.80–0.07) N = 10 | 3.60 ± 1.20 (1.80–5.40) N = 23 |
| | O | 1.20 ± 0.60 (0.60 –2.40) N = 7 | 1.20 ± 1.20 (0.60–3.00) N = 10 | 1.80 ± 0.60 (0.60 –3.00) N = 16 |
| Peak frequency | RC | 13.59 ± 0.39 (12.92–14.08) N = 10 | 13.41 ± 0.42 (12.74–14.11) N = 12 | 13.64 ± 0.47 (12.68–14.36) N = 25 |
| | LC | 13.63 ± 0.37 (12.90–14.31) N = 11 | 13.46 ± 0.36 (12.65–13.90) N = 11 | 13.58 ± 0.35 (12.73–14.15) N = 23 |
| | F | 13.76 ± 0.35 (13.01–14.34) N = 11 | 13.52 ± 0.50 (12.56–14.24) N = 11 | 13.63 ± 0.47 (12.63–14.31) N = 23 |
| | O | 13.69 ± 0.42 (13.10–14.40) N = 8 | 13.78 ± 0.56 (13.17–14.81) N = 11 | 13.60 ± 0.58 (12.46–14.45) N = 17 |
| Amplitude | RC | 23.18 ± 4.34 (15.99–29.42) N = 9 | 20.66 ± 5.34 (12.11–29.55) N = 12 | 21.78 ± 6.72 (10.25–35.78) N = 25 |
| | LC | 19.83 ± 6.60 (7.14–29.69) N = 11 | 21.23 ± 7.59 (10.26–35.43) N = 12 | 22.69 ± 7.29 (10.77–38.05) N = 23 |
| | F | 20.17 ± 5.32 (10.83–26.75) N = 12 | 16.02 ± 3.37 (12.18–22.65) N = 10 | 19.98 ± 5.83 (11.82–30.26) N = 24 |
| | O | 13.31 ± 1.99 (10.75–16.73) N = 7 | 13.56 ± 2.06 (9.82–17.20) N = 11 | 15.45 ± 4.37 (10.50–25.17) N = 16 |

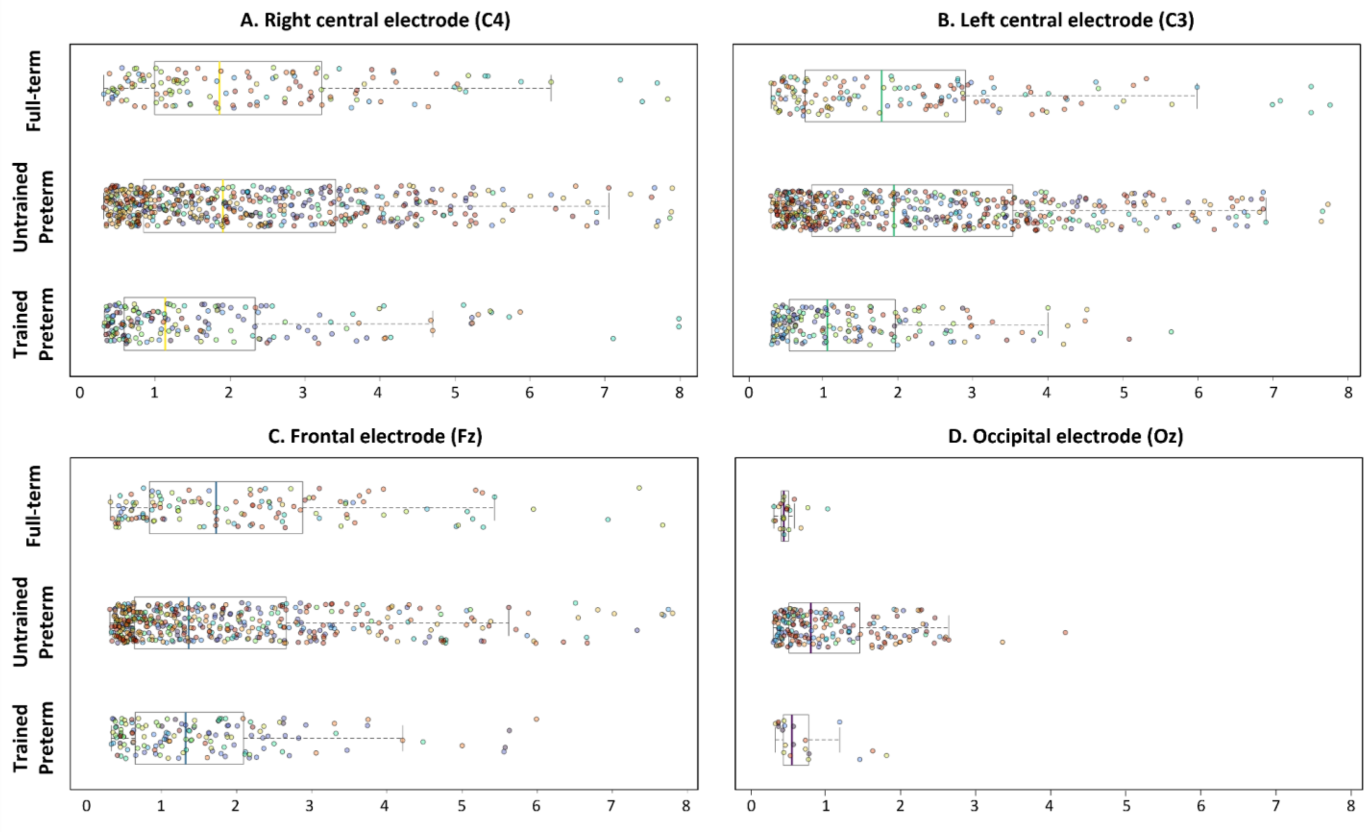


**Supplementary information. Figure 5.** Event-level distribution of sleep spindle durations across groups at the representative electrode of each cluster

Quantification of sleep spindle features was performed within each group (full-term, trained preterm, and untrained preterm infants) at the representative electrode of each cluster (C4, C3, Fz, Oz). Each point represents an individual sleep spindle, with all spindles from each infant included to capture within-subject variability. Boxplots summarise group-level distributions at each representative electrode. Colors indicate individual subjects, with all spindles from the same subject sharing the same color.

**A.** C4: full-term: mean = 2.31s ± 1.71s (range = 0.31s – 7.83s), untrained preterm: mean = 2.30s ± 1.71s (range = 0.31s – 7.90s), trained preterm: mean = 1.72s ± 1.58s (range = 0.32s – 7.98s). **B.** C3: full-term: mean = 2.16s ± 1.69s (range = 0.32s – 7.76s), untrained preterm: mean = 2.38s ± 1.75s (range = 0.31s – 7.73s), trained preterm: mean = 1.42s ± 1.09s (range = 0.32s – 5.65s). **C.** Fz: full-term: mean = 2.10s ± 1.58s (range = 0.32s – 7.67s), untrained preterm: mean = 1.93 s ± 1.60s (range = 0.31s – 7.80s), trained preterm: mean = 1.57s ± 1.17s (range = 0.33s – 5.99s). **D.** Oz: full-term: mean = 0.49 s ± 0.16s (range = 0.32s – 1.03s), untrained preterm: mean = 1.06 s ± 0.70s (range = 0.31s – 4.20s), trained preterm: mean = 0.73s ± 0.44s (range = 0.34s – 1.82s).

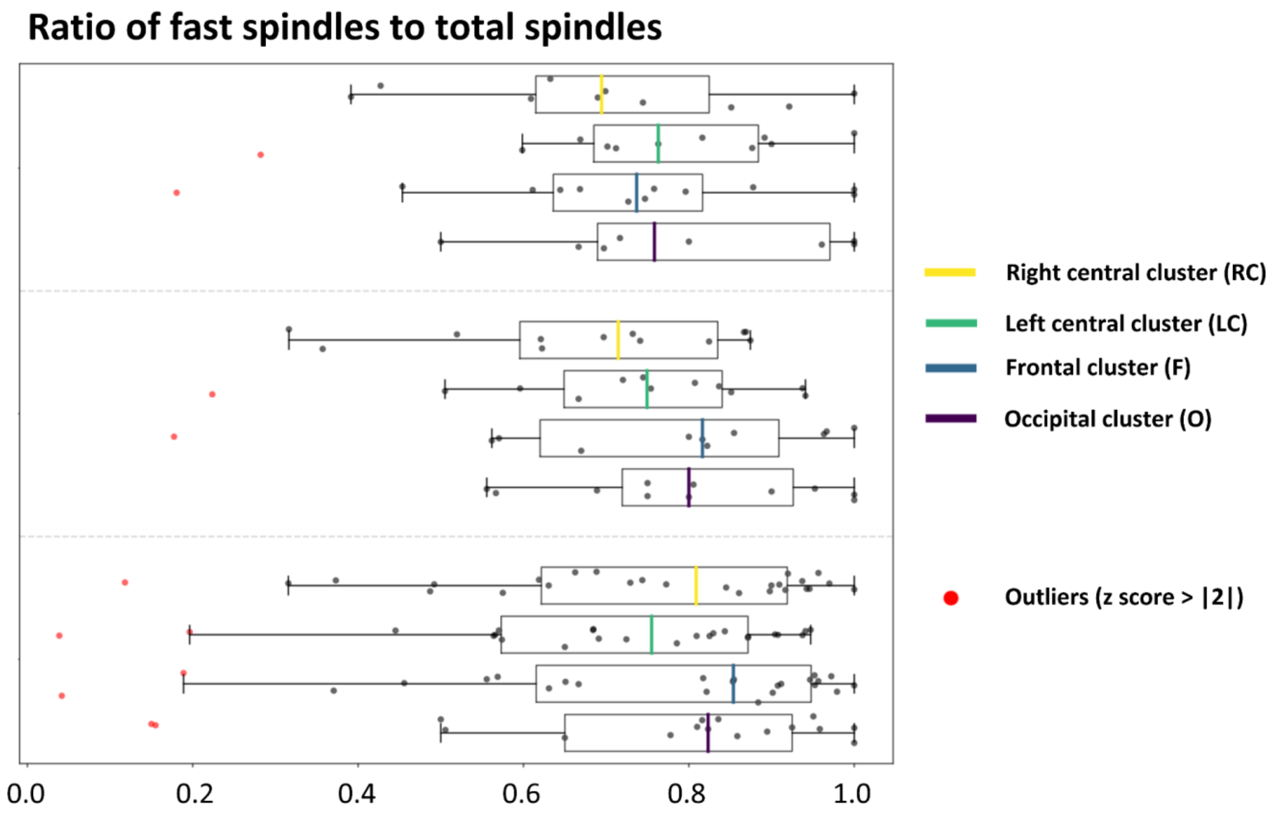


**Supplementary information. Figure 6.** Proportion of fast (13-16Hz) VS slow spindles (11-13Hz) in the three groups

Quantification of the ratio of fast spindles to total spindles and identification of outliers (z-score ≥ |2|) within each group (full-term, trained preterm, and untrained preterm infants) and for each electrode cluster (right-central, left-central, occipital, and frontal). Each boxplot represents the mean value per infant within each group and cluster for all analysed spindle features.

**A**. RC: full-term: mean = 0.79 ± 0.18 (range = 0.39s – 1.00), untrained preterm: mean = 0.74 ± 0.23 (range = 0.12-1.00), trained preterm: mean = 0.67 ± 0.19 (range = 0.32 – 0.87) / LC: full-term: mean = 0.75 ± 0.19 (range = 0.28s – 1.00), untrained preterm: mean = 0.70 ± 0.23 (range = 0.04-1.00), trained preterm: mean = 0.72 ± 0.20 (range = 0.22 – 0.94) / F: full-term: mean = 0.71 ± 0.22 (range = 0.18s – 1.00), untrained preterm: mean = 0.74 ± 0.26 (range = 0.04-1.00), trained preterm: mean = 0.75 ± 0.24 (range = 0.18 – 1.00). / O: full-term: mean = 0.79 ± 0.18 (range = 0.50s – 1.00), untrained preterm: mean = 0.74 ± 0.27 (range = 0.15-1.00), trained preterm: mean = 0.80 ± 0.16 (range = 0.56 – 1.00).

**Table 4. Summary of LMM results for group, cluster, and interaction (group x cluster) effects across sleep spindle features**

| Effect | Sum Sq | Mean Sq | NumDF | DenDF | F value | p value |
|---|---|---|---|---|---|---|
| | | | **ISI** | | | |
| **Group** | 48.90 | 24.45 | 2.00 | 37.68 | 0.06 | 0.94 |
| **Cluster** | 1428.00 | 476.01 | 3.00 | 62.74 | 1.24 | 0.30 |
| **Group:Cluster** | 3176.90 | 529.48 | 6.00 | 62.43 | 1.38 | 0.24 |
| *marginal R2 = 0.07. conditional R2 = 0.39* | | | | | | |
| | | | **ISI** | | | |
| **Group** | 0.30 | 0.15 | 2.00 | 42.98 | 0.07 | 0.93 |
| **Cluster** | 2.45 | 0.82 | 3.00 | 114.08 | 0.38 | 0.77 |
| **Group:Cluster** | 27.00 | 4.50 | 6.00 | 114.11 | 2.09 | 0.06 |
| *marginal R2 = 0.05. conditional R2 = 0.46* | | | | | | |
| | | | **Duration** | | | |
| **Group** | 0.41 | 0.21 | 2.00 | 44.13 | 0.84 | 0.44 |
| **Cluster** | 31.84 | 10.61 | 3.00 | 111.59 | 43.26 | <10e-15 |
| **Group:Cluster** | 2.70 | 0.45 | 6.00 | 111.58 | 1.83 | 0.10 |
| *marginal R2 = 0.31. conditional R2 = 0.68* | | | | | | |
| | | | **Density** | | | |
| **Group** | 0.00 | 0.00 | 2.00 | 45.35 | 3.38 | 0.04 |
| **Cluster** | 0.02 | 0.01 | 3.00 | 113.77 | 64.36 | <10e-15 |
| **Group:Cluster** | 0.00 | 0.00 | 6.00 | 113.72 | 1.06 | 0.39 |
| *marginal R2 = 0.42. conditional R2 = 0.72* | | | | | | |
| | | | **Peak frequency** | | | |
| **Group** | 0.02 | 0.01 | 2.00 | 46.64 | 0.19 | 0.82 |
| **Cluster** | 0.44 | 0.15 | 3.00 | 117.12 | 2.96 | 0.04 |
| **Group:Cluster** | 0.66 | 0.11 | 6.00 | 117.07 | 2.24 | 0.04 |
| *marginal R2 = 0.04. conditional R2 = 0.76* | | | | | | |
| | | | **Amplitude** | | | |
| **Group** | 11.66 | 5.83 | 2.00 | 45.22 | 0.53 | 0.59 |
| **Cluster** | 1057.22 | 352.41 | 3.00 | 113.15 | 32.04 | < 10e-16 |
| **Group:Cluster** | 102.73 | 17.12 | 6.00 | 113.11 | 1.56 | 0.17 |
| *marginal R2 = 0.19. conditional R2 = 0.75* | | | | | | |

**Table 5. Summary of post hoc statistics (emmeans) in the comparison of duration between clusters**

**RC:** Right-central cluster, **LC:** Left-central cluster, **F:** Frontal cluster, **O:** Occipital cluster

| Duration | | | | | |
|---|---|---|---|---|---|
| **Contrast** | **Estimate** | **SE** | **df** | **t ratio** | **p value FDR corrected** |
| | | **Effect of the factor cluster** | | | |
| **RC-LC** | 0.11 | 0.11 | 111.00 | 0.99 | 0.33 |
| **RC- F** | 0.45 | 0.11 | 114.00 | 3.95 | 2e-4 |
| **RC- O** | 1.29 | 0.12 | 115.00 | 10.50 | <10–4 |
| **LC- F** | 0.34 | 0.11 | 113.00 | 3.02 | 2e-3 |
| **LC - O** | 1.18 | 0.12 | 114.00 | 9.70 | <10–4 |
| **F - O** | 0.84 | 0.12 | 115.00 | 6.74 | <10–4 |

**Table 6. Summary of post hoc statistics (emmeans) in the comparison of density between groups and clusters**

**RC:** Right-central cluster, **LC:** Left-central cluster, **F:** Frontal cluster, **O:** Occipital cluster

**FT**: Full-term infants, **UP**: Untrained preterm infants, **TP**: Trained preterm infants

| Density | | | | | |
|---|---|---|---|---|---|
| **Contrast** | **Estimate** | **SE** | **df** | **t ratio** | **p value FDR corrected** |
| | | **Effect of the factor group** | | | |
| **FT - UP** | -0.01 | 4e-3 | 47.10 | -2.45 | 0.05 |
| **FT - TP** | -4e-3 | 0.01 | 46.10 | -0.76 | 0.45 |
| **TP - UP** | -0.01 | 4e-3 | 45.10 | -1.60 | 0.17 |
| | | **Effect of the factor cluster** | | | |
| **RC-LC** | -2e-4 | 3e-3 | 113.00 | 0.06 | 0.95 |
| **RC- F** | 3e-3 | 3e-3 | 114.00 | 1.45 | 0.18 |
| **RC- O** | 0.03 | 3e-3 | 115.00 | 12.17 | <10–4 |
| **LC- F** | 3e-3 | 3e-3 | 115.00 | 1.50 | 0.18 |
| **LC - O** | 0.03 | 3e-3 | 115.00 | 12.18 | <10–4 |
| **F - O** | 0.03 | 3e-3 | 116.00 | 10.75 | <10–4 |

**Table 7. Summary of post hoc statistics (emmeans) in the comparison of peak frequency between clusters and groups x clusters**

**RC:** Right-central cluster, **LC:** Left-central cluster, **F:** Frontal cluster, **O:** Occipital cluster
**FT**: Full-term infants, **UP**: Untrained preterm infants, **TP**: Trained preterm infants

**Peak frequency**

| Contrast | Estimate | SE | df | t ratio | p value FDR corrected |
|---|---|---|---|---|---|
| **Effect of the factor cluster** | | | | | |
| **RC-LC** | -0.02 | 0.05 | 116.00 | -0.36 | 0.72 |
| **RC- F** | -0.07 | 0.05 | 117.00 | -1.34 | 0.28 |
| **RC- O** | -0.15 | 0.05 | 117.00 | -2.75 | 0.04 |
| **LC- F** | -0.05 | 0.05 | 117.00 | -0.99 | 0.39 |
| **LC - O** | -0.13 | 0.05 | 116.00 | -2.42 | 0.05 |
| **F - O** | -0.08 | 0.05 | 117.00 | -1.48 | 0.28 |
| **Effect of the factor group x cluster** | | | | | |
| **FT** | | | | | |
| **RC-LC** | -0.04 | 0.10 | 115.00 | -0.36 | 0.86 |
| **RC- F** | -0.11 | 0.10 | 118.00 | -1.06 | 0.70 |
| **RC- O** | -0.12 | 0.11 | 117.00 | -1.10 | 0.70 |
| **LC- F** | -0.07 | 0.10 | 118.00 | -0.74 | 0.70 |
| **LC - O** | -0.09 | 0.11 | 116.00 | -0.81 | 0.70 |
| **F - O** | -0.01 | 0.11 | 119.00 | -0.13 | 0.90 |
| **UP** | | | | | |
| **RC-LC** | 0.08 | 0.06 | 116.00 | 1.22 | 0.79 |
| **RC- F** | 0.02 | 0.07 | 116.00 | 0.23 | 0.82 |
| **RC- O** | 0.05 | 0.07 | 117.00 | 0.65 | 0.79 |
| **LC- F** | -0.06 | 0.07 | 116.00 | -0.97 | 0.79 |
| **LC - O** | -0.03 | 0.07 | 117.00 | -0.44 | 0.79 |
| **F - O** | 0.03 | 0.07 | 116.00 | 0.44 | 0.79 |
| **TP** | | | | | |
| **RC-LC** | -0.10 | 0.09 | 115.00 | -1.05 | 0.36 |
| **RC- F** | -0.11 | 0.09 | 116.00 | -1.19 | 0.35 |
| **RC- O** | -0.37 | 0.09 | 116.00 | -3.94 | 10–4 |
| **LC- F** | -0.01 | 0.10 | 116.00 | -0.14 | 0.89 |
| **LC - O** | -0.27 | 0.10 | 116.00 | -2.81 | 0.02 |
| **F - O** | -0.26 | 0.09 | 115.00 | -2.72 | 0.02 |

**Table 8. Summary of post hoc statistics (emmeans) in the comparison of amplitude between cluster**

**RC:** Right-central cluster, **LC:** Left-central cluster, **F:** Frontal cluster, **O:** Occipital cluster

**Amplitude**

| Contrast | Estimate | SE | df | t ratio | p value FDR corrected |
|---|---|---|---|---|---|
| **Effect of the factor cluster** | | | | | |
| **RC-LC** | 0.37 | 0.75 | 114.00 | 0.49 | 0.62 |
| **RC- F** | 2.54 | 0.77 | 115.00 | 3.32 | 2e-3 |
| **RC- O** | 7.34 | 0.83 | 115.00 | 8.84 | <10–4 |
| **LC- F** | 2.17 | 0.75 | 115.00 | 2.91 | 0.01 |
| **LC - O** | 6.97 | 0.82 | 115.00 | 8.55 | <10–4 |
| **F - O** | 4.80 | 0.82 | 116.00 | 5.84 | <10–4 |

#### Slow oscillation-spindle coupling in infancy

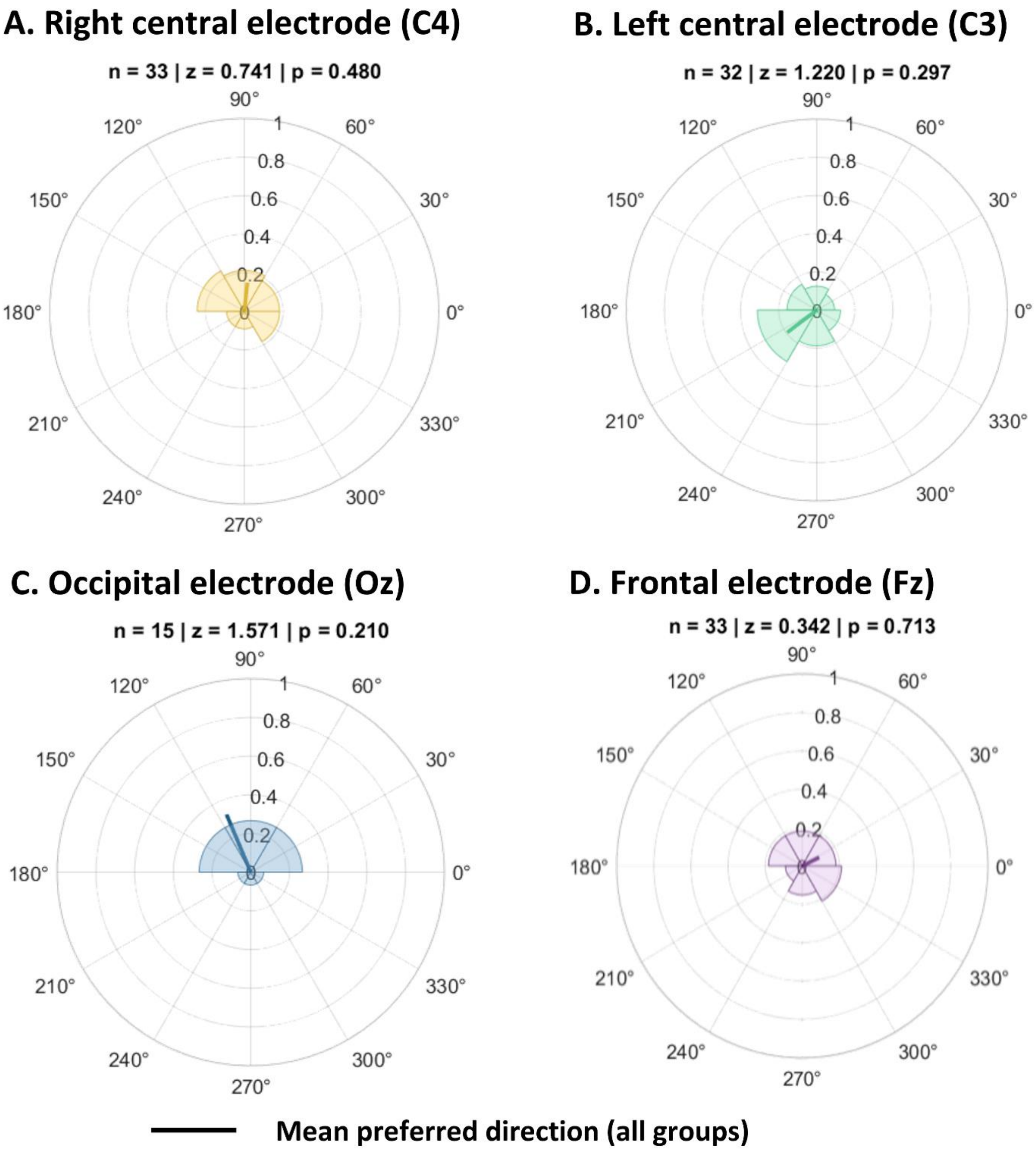


**Supplementary Information. Figure 7. Slow oscillation–spindle coupling**

Preferred phase (in radians) and resultant vector length of slow oscillations at the onset of spindle recorded at C4. Analyses were performed in an exploratory way with data from our three groups: full-term, untrained preterm and trained preterm infants.

Results are shown separately at the representative electrode of each cluster: **A.** C4, **B.** C3, **C.** Fz, **D.** Oz. Probability density distributions of phase angles (histograms) are also displayed for each electrode. Phase uniformity was assessed with Rayleigh tests (z). No significant deviation from a uniform distribution was shown for any electrode.

Slow-oscillations detection

As sleep spindles, slow-oscillations (SO) were detected using an automated algorithm originally developed for adults [1] and subsequently adapted to the specific characteristics of infants.The first step consisted of band-pass filtering the signal using a finite impulse response (FIR) filter tuned to the SO frequency range (0.5–1.25 Hz). After filtering, the signal was segmented into multiple events corresponding to potential SO. Zero crossings were identified, and each interval between two consecutive positive zero crossings was defined as a candidate event. A duration criterion was then applied, consistent with thresholds established for both adults [1] and 2–3-month-old infants [2], retaining only events lasting between 0.8 and 2 seconds. Events falling outside this range were discarded. Finally, the peak-to-peak amplitude of each candidate SO was computed to establish a relative amplitude threshold based on the 75th percentile of the amplitude distribution. Only events exceeding this relative threshold were classified as SO.

#### Coupling

Once SO and sleep spindles had been detected, we examined their coupling using a method similar to that described by [1]. Signals were first band-pass filtered using linear-phase FIR filters. The SO band was defined as 0.5–1.25 Hz, and the spindle band as 11–16 Hz. Co-occurring slow oscillation–spindle events were defined as the events of interest for subsequent analyses and were identified when spindle onset occurred within the temporal window of the SO. Hilbert transforms were applied to both SO and spindle signals. This enabled extraction of the instantaneous phase of the SO (angle of the Hilbert-transformed SO signal) and the instantaneous amplitude of the spindle band (absolute value of the Hilbert-transformed spindle signal). We then assessed phase–amplitude SO–spindle coupling by extracting the SO phase at the time the spindle reached its highest amplitude.

However, this technique yielded an insufficient number of co-occurring SO-spindle events for reliable analysis (full-term, untrained preterm, and trained preterm infants, respectively: N = 3, 11, 7 in C4; N = 2, 6, 1 in Oz; N = 4, 11, 6 in C3; and N = 5, 15, 4 in Fz). Consequently, we decided to focus on spindle onset rather than spindle amplitude. Since the maximum spindle duration was set to 8s (compared to 2s in adults, [1] ; and 3s in 2–3-month-old infants,[2]) this may have reduced the probability that the peak spindle amplitude would occur within the SO time window. Instead of assessing phase–amplitude SO–spindle coupling, we then identified spindle onsets occurring after SO onset and extracted the corresponding SO phase at that time point. Phase-onset coupling was then quantified using circular statistics. The preferred phase was computed as the circular mean of SO phases at spindle onset, and coupling strength was quantified as the length of the mean resultant vector, which reflects how consistently spindle onsets occur at a given SO phase.

Due to the low number of co-occurring so-spindle events and the short recording duration available, we chose to conduct an exclusively exploratory analysis. In this context, rather than analysing our three groups separately, we pooled them together in order to investigate a potential coupling at 2 months of corrected age in a global manner. Statistical analyses were performed using the CircStats toolbox [3]. In a first step, the distributions of preferred phases were modeled using the von Mises distribution in order to estimate the mean direction and the concentration parameter, thereby assessing whether a well-defined and sufficiently concentrated preferred direction was present. Rayleigh tests were then applied to determine whether the observed phase distributions deviated from uniformity by exhibiting a significant clustering around a specific direction. When the Rayleigh test was significant, Watson–Williams tests were subsequently used to compare mean directions between groups, under the assumption of a unimodal distribution consistent with the estimated von Mises parameters.

As shown in Supplementary Information. Figure 7, no significant coupling between slow oscillations and sleep spindles was evidenced.
